# Neurovirulence of the Australian outbreak Japanese Encephalitis virus genotype 4 is lower compared to genotypes 2 and 3 in mice and human cortical brain organoids

**DOI:** 10.1101/2023.04.26.538504

**Authors:** Wilson Nguyen, Narayan Gyawali, Romal Stewart, Bing Tang, Abigail L. Cox, Kexin Yan, Thibaut Larcher, Cameron Bishop, Nicholas Wood, Gregor J. Devine, Andreas Suhrbier, Daniel J. Rawle

**Author notes:** should be considered joint last authors (Andreas Suhrbier, Daniel J. Rawle).

## Abstract

**Background:** Human infections with Japanese encephalitis virus (JEV) are a leading cause of viral encephalitis. An unprecedented outbreak of JEV genotype 4 was recently reported in Australia, with an isolate (JEV_NSW/22_) obtained from a stillborn piglet brain.

**Methods:** Herein we compared the neuropathology of JEV_NSW/22_, JEV_FU_ (genotype 2) and JEV_Nakayama_ (genotype 3) in adult C57BL/6J wild-type mice, mice deficient in interferon regulatory factor 7 (*Irf7^-/-^*), and mice deficient in type I interferon receptor (*Ifnar^-/-^*), as well as in human cortical brain organoids (hBOs). Using human serum post-Imojev vaccination, we performed neutralisation assays to determine JEV_NSW/22_ susceptibility to vaccine responses.

**Findings:** In C57BL/6J and *Irf7^-/-^* mice with lethal outcomes, brain infection and histopathological lesions recapitulated those seen in humans and primates. JEV was universally lethal in *Ifnar^-/-^* mice by day 3 with histological signs of brain hemorrhage, but produced no other detectable brain infection or lesions, with viral protein detected in blood vessels but not neurons. We thus describe a new *Irf7^-/-^* mouse model for JEV_NSW/22_, which had increased viremia compared to C57BL/6J mice, allowing for lethal neuroinvasive infection in one mouse. Overall, JEV_NSW/22_ was less neurovirulent than other JEV isolates in C57BL/6J and *Irf7^-/-^* mice, and was more sensitive to type I interferon. All JEV isolates showed robust cytopathic infection of human cortical brain organoids, albeit lower for JEV_NSW/22_. We also show that Imojev vaccination in humans induced neutralizing antibodies against JEV_NSW/22_, with the level of cross-neutralisation related to the conservation in envelope protein amino acid sequences for each isolate.

**Interpretation:** Our study establishes JEV_NSW/22_ mouse models of infection, allowing for possible lethal neuroinvasive infection that was rarer than for other JEV genotypes. JEV vaccination regimens may afford protection against this newly emerged JEV genotype 4 strain, although neutralizing antibody responses are sub-optimal.

**Funding:** QIMRB received a generous philanthropic donation from the Brazil Family Foundation awarded to D.J.R. to support Japanese Encephalitis virus research at QIMRB. A.S. holds an Investigator grant from the National Health and Medical Research Council (NHMRC) of Australia (APP1173880). We also acknowledge the intramural grant from QIMR Berghofer awarded to R.S. and D.J.R. for purchase of the CelVivo Clinostar incubator for producing human cortical brain organoids. The project “Japanese encephalitis vaccine via the intradermal route in children and adults (JEVID-2): A clinical trial comparing the immunogenicity and safety of Japanese encephalitis vaccine administered by subcutaneous and intradermal routes” being conducted by G.D., N.G., and N.W. was funded by the Sydney Children’s Hospitals Network and New South Wales Health.

**Research in context:** *Evidence before the study:* JEV from the historically rare genotype 4 recently emerged in Australia, causing an unprecedented outbreak, with 44 human cases and 7 fatalities. While a range of JEV mouse models have been reported, none of them infect adult mice with a genotype 4 isolate. The efficacy of current vaccines for this JEV genotype are also unclear.

*Added value of this study:* We establish well characterised adult and subcutaneously infected mouse models for JEV which recapitulate many aspects of human disease including lethal neuroinvasive infection and severe histopathological lesions. Prolonged viremia was significantly associated with lethal neuroinvasiveness in *Irf7^-/-^*mice. We demonstrate that a genotype 4 Australian isolate, JEV_NSW/22_, exhibited markedly diminished lethal neuroinvasion compared to other JEV genotypes. Using serum from Imojev vaccine recipients, neutralizing antibodies against JEV_NSW/22_ were present, albeit at sub-optimal titers.

*Implications of all the available evidence:* The establishment of well characterised adult mouse models of JEV_NSW/22_ with rare neuropenetrance after peripheral inoculation that recapitulate human disease is an important tool that can now be deployed in pre-clinical studies and to understand disease pathogenesis. Our study suggests that new vaccines should be developed against circulating JEV strains for optimal neutralizing antibody responses.

## Introduction

Japanese Encephalitis virus (JEV) is a single-stranded positive-sense RNA virus from the *Flaviviridae* family that is transmitted from amplifying hosts (primarily pigs and wading birds) via mosquitoes to humans (1). While infection is usually asymptomatic, encephalitis can develop in ≈1 in 250 people, with ∼30% of encephalitic cases becoming fatal, and 30-50% of non-fatal encephalitic cases retaining persistent neurological symptoms including seizures, speech impediments and paralysis (2, 3). JEV is the leading cause of viral encephalitis in Asia, with ∼70,000 cases and ∼20,000 deaths per annum (4). After a bite from an infected mosquito, JEV replicates in peripheral blood monocytes producing a viremia that, in some cases, leads to virus crossing the blood brain barrier (5). JEV primarily infects neurons in the brain (6) leading to uncontrolled inflammation (encephalitis) and neuronal cell death (2). There are several available and effective vaccines against JEV (7), but there are no specific licensed treatments.

JEV exists as five genotypes (1 to 5) which are phylogenetically, antigenically and geographically distinct (8–10). Genotype 3 was the main genotype endemic in Asia until 1990, after which genotype I has dominated since (11). Genotype 2 has been identified in Malaysia, Indonesia, Papua New Guinea and Australian outbreaks from 1970-2000. Genotype 5 has been identified in China, and is now dominant in South Korea (12). Genotype 4 was the least common genotype worldwide, having only been identified in mosquitoes from Indonesia and Papua New Guinea (13). In February 2021, a fatal JEV infection occurred on the Tiwi Islands, 80 km off the coast of Darwin, Australia (14). Sequencing identified that the virus belonged to the historically rare JEV genotype 4 (14). In 2022, a geographically widespread outbreak throughout most Australian states was attributed to JEV genotype 4, which caused 44 confirmed human cases and 7 deaths (15). Outbreaks occurred in several piggeries, causing high abortion and stillbirth rates in sows (16). Based on proximity to piggeries, estimates indicate that ∼740,000 people may be at risk of being infected by JEV in Australia (16). Given that *Culex annulirostris* mosquitoes, considered the primary vector for JEV in Australia (15, 16), and amplifying vertebrate hosts including wading birds and pigs are widespread, it is possible that JEV may become, or is already, endemic in Australia (15). Murray Valley Encephalitis virus (MVEV) and Kunjin virus are phylogenetically closely related to JEV and have similar vector and reservoir hosts, and are endemic to Australia (17).

Efforts to develop treatments for JEV neuropathology are hindered by the difficulties in diagnosing early infection (13) before the virus has infected central nervous system (CNS) neurons; and once the virus has infected CNS neurons, treatments must also be able to effectively enter the brain. While a range of JEV mouse models have been reported (Supplementary Table 1) (18), there are no studies of genotype 4 in adult mice, with the only study on genotype 4 JEV using 3-week-old mice (19). Well-characterized, adult mouse models of recent genotype 4 isolates would represent useful tools to study mechanisms of infection and disease and evaluate potential new interventions. Herein, we compare and contrast a JEV genotype 4 isolate from the recent Australian outbreak (JEV_NSW/22_) with historical JEV isolates (genotypes 2 and 3) in three mouse strains, human cortical brain organoids and *in vitro* neutralisation assays with human serum after JEV vaccination.

## Materials and methods

### Ethics statement and regulatory compliance

All mouse work was conducted in accordance with the “Australian code for the care and use of animals for scientific purposes” as defined by the National Health and Medical Research Council of Australia. Mouse work was approved by the QIMR Berghofer Medical Research Institute (MRI) Animal Ethics Committee (P3746, A2108-612). Mice were euthanized using CO_2_.

Breeding and use of GM mice was approved under a Notifiable Low Risk Dealing (NLRD) Identifier: NLRD_Suhrbier_Oct2020: NLRD 1.1(a). Use of Imojev was approved under NLRD_Suhrbier_Oct2019: NLRD2.1(d).

All infectious JEV work was conducted in a dedicated suite in a biosafety level-3 (PC3) facility at the QIMR Berghofer MRI (Australian Department of Agriculture, Water and the Environment certification Q2326 and Office of the Gene Technology Regulator certification 3445). All work was approved by the QIMR Berghofer MRI Safety Committee (P3746).

Human serum samples before and after Imojev vaccination were collected from 9 participants with human research ethics approval from Sydney Children’s Hospitals Network Human Research Ethics Committee (2022/ETH02471 HCHNHREC) for an ongoing project led by Dr. Nicholas Wood; “Japanese encephalitis vaccine via intradermal route in children and adults (JEVID-2): Critical policy relevant research for Australia”.

Human serum samples after Imojev vaccination were also collected from 10 participants with approval from QIMR Berghofer MRI Human Research Ethics Committee (P3476) provided for neutralisation assays (including MVEV cross-neutralisation).

Research with JEV were approved under the Queensland Biosecurity Act, Scientific Research Permit (restricted matter) – Permit number PRID000916.

### Cell lines and culture

Vero E6 (C1008, ECACC, Wiltshire, England; obtained via Sigma Aldrich, St. Louis, MO, USA), BHK-21 (ATCC# CCL-10), and C6/36 cells (ATCC# CRL-1660) were cultured in medium comprising RPMI 1640 (Gibco), supplemented with 10% fetal bovine serum (FBS), penicillin (100LIU/ml)/streptomycin (100 μg/ml) (Gibco/Life Technologies) and L-glutamine (2 mM) (Life Technologies). Cells were cultured at 37°C and 5% CO_2_. Cells were routinely checked for mycoplasma (MycoAlert Mycoplasma Detection Kit MycoAlert, Lonza) and FBS was assayed for endotoxin contamination before purchase (20).

Mouse embryonic fibroblasts (MEFs) were either wild type or *Irf3/7^-/-^*and have been described previously (21). MEFs were cultured in Dulbecco’s Modified Eagle Medium (DMEM) (Gibco), supplemented with 50μg/ml Penicillin/50μg/ml Streptomycin and 10% Fetal Bovine Serum (FBS). Cells were cultured at 37°C and 5% CO_2_. For virus growth kinetics, MEFs were seeded at 2×10^5^ cells/ml in 12 or 24 well plates one day prior to infection at multiplicity of infection (MOI) 0.1 of the indicated JEV or MVEV. After 1 hr incubation, MEFs were washed twice with 1 ml PBS, and culture medium was added and sampled daily for virus titrations by CCID_50_ (see below).

RENcell VM Human Neural Progenitor Cell Line (Sigma Aldrich) were cultured in medium comprising DMEM F-12 (Thermo Fisher Scientific), penicillin (100LIU/ml)/streptomycin (100 μg/ml) (Gibco/Life Technologies), 20 ng/ml FGF (STEMCELL Technologies), 20 ng/ml EGF (STEMCELL Technologies), and 2% B27 supplements (Thermo Fisher Scientific). Cells were detached using StemPro Accutase Cell Dissociation Reagent (Thermo Fisher Scientific), and were cultured in Matrigel (Sigma). For crystal violet staining of remaining cells after infection, formaldehyde (7.5% w/v)/crystal violet (0.05% w/v) was added to wells overnight, plates washed twice in water, and plates dried overnight.

### Virus isolates and culture

JEV_Nakayama_ (GenBank: EF571853), JEV_FU_ (GenBank: AF217620), and Imojev were obtained from A/Prof. Gregor Devine (QIMR Berghofer MRI). JEV_NSW/22_ (GenBank: OP904182) was a gift from Dr. Peter Kirkland (Elizabeth Macarthur Agriculture Institute, New South Wales, Australia). MVEV_TC123130_ (GenBank: JN119814.1) was obtained from the “Doherty Virus Collection” currently held at QIMR Berghofer MRI. Virus stocks were generated by infection of C6/36 cells (for JEV and MVEV) or Vero E6 cells (for Imojev and YFV 17D) at multiplicity of infection (MOI) ≈0.1, with supernatant collected after ∼5 days, cell debris removed by centrifugation at 3000 x g for 15 min at 4°C, and virus aliquoted and stored at –80°C. Virus stocks used in these experiments underwent less than three passages in our laboratory, with prior passage history in Supplementary Figure 1. Virus titers were determined using standard CCID_50_ assays (see below).

### Validation of virus stock sequences

Viral RNA was extracted from virus stock culture supernatants using NucleoSpin RNA Virus kit (Machery Nagel) as per manufacturer’s instructions. cDNA was synthesized using ProtoScipt First Strand cDNA Synthesis Kit (New England Biolabs) as per manufacturer’s instructions.

PCR was performed using Q5 High-Fidelity 2X Master Mix (New England Biolabs) as per manufacturer’s instructions with the following primers; JEV envelope Forward 5’ GGAAGCATTGACACATGTGC 3’ and Reverse 5’ TCTGTGCACATACCATAGGTTGTG 3’, and MVEV envelope Forward 5’ GAGCATTGACACATGCGCAAAG and Reverse 5’ TGTGCACATCCCATAAGTGGTTC 3’. PCR products were run on a 1% agarose gel and DNA was extracted using Monarch DNA Gel Extraction Kit (New England Biolabs). DNA was sequenced by Sanger sequencing using either the forward or reverse primer. Sequences of our virus stocks matched the sequences on GenBank (Supplementary Figure 1).

### CCID_50_ assays

Vero E6 cells were plated into 96 well flat bottom plates at 2×10^4^ cells per well in 100 µl of medium. For tissue titrations, tissues were homogenized in tubes each containing 4 ceramic beads twice at 6000 x g for 15 seconds, followed by centrifugation twice at 21,000 x g for 5 min before 5 fold serial dilutions in 100 µl RPMI 1640 supplemented with 2% FBS. For cell culture supernatant or mouse serum, 10 fold serial dilutions were performed in 100 µl RPMI 1640 supplemented with 2% FBS. A volume of 100 µl of serially diluted samples were added to each well of 96 well plate containing Vero E6 cells and the plates cultured for 5-7 days at 37°C and 5% CO_2_. The virus titer was determined by the method of Spearman and Karber.

### Mouse infections

C57BL/6J mice were purchased from the Animal Resources Center, Canning Vale WA, Australia. *Ifnar^-/-^*were kindly provided by Dr P. Hertzog (Monash University, Melbourne, Australia). *Irf7^-/-^* mice were kindly provided by T. Taniguchi (University of Tokyo) (22–24). C57BL/6N mice were purchased from The Jackson Laboratory (stock no. 005304).

C57BL/6NJ^Δ*Nnt8-12*^ were generated by The Australian Phenomics Network, Monash University, Melbourne, Australia as described (25). Mice used in this study were female, except *Irf7^-/-^* mice where both males and females were used. The age/age range at infection is indicated in the figure legends. Mice were sorted into groups so that each group had a similar mean age and age distribution, and in the case of *Irf7^-/-^* mice, had equal numbers of males and females in each group. All mice strains except C57BL/6J were bred in-house at QIMR Berghofer MRI, and mice housing conditions were described previously (26). Mice were infected subcutaneously (s.c.) at the base of the tail with 100 µl of virus inoculum (doses ranging from 5×10^2^ to 5×10^5^ as indicated in the figure legends). Serum was collected via tail nicks in to Microvette Serum-Gel 500 µl blood collection tubes (Sarstedt, Numbrecht, Germany). Mice were weighed and monitored for disease manifestations and were humanely euthanized using CO_2_ based on a score card system (Supplementary Figure 2B). At necropsy, brain and spleens were collected for virus titrations by CCID_50_ assays and/or for histology.

### Histopathology and immunohistochemistry

Brains and spleens were fixed in 10% formalin and embedded in paraffin. Human cortical brain organoids were embedded in agarose by adding 4% melted agarose and incubating on ice to solidify, prior to standard paraffin embedding. Sections were stained with H&E (Sigma Aldrich) and slides were scanned using Aperio AT Turbo (Aperio, Vista, CA USA) and analyzed using Aperio ImageScope software (LeicaBiosystems, Mt Waverley, Australia) (v10). Leukocyte infiltrates were quantified by measuring nuclear (strong purple staining) / cytoplasmic (total red staining) pixel ratios in scanned H&E stained images, and was undertaken using Aperio Positive Pixel Count Algorithm (Leica Biosystems) (27).

For anti-flavivirus non-structural protein 1 (NS1) immunohistochemistry using 4G4, sections were affixed to positively charged adhesive slides and air-dried overnight at 37°C. Sections were dewaxed and rehydrated through xylol and descending graded alcohols to water. Sections were transferred to Dako Epitope Retrieval Solution (pH 9.0) and subjected to heat antigen retrieval (100°C for 20 min) using the Biocare Medical de-cloaking chamber, and slides allowed to cool for 20 minutes before transferring to TBS plus 0.025% Tween-20 (TBS-T). Endogenous mouse Ig was blocked by incubating sections with donkey anti-mouse Fab fragments (Jackson Immunoresearch) diluted 1:50 in Biocare Medical Rodent block M for 60 minutes. Sections were washed three times in TBS-T, then incubated with anti-mouse Fc for 15 minutes, before a further three washes in TBS-T. Nonspecific antibody binding was inhibited by incubation with Biocare Medical Background Sniper with 1% BSA and 20% donkey serum and 20% goat serum for 15 minutes. Primary antibody 4G4 (mouse anti-flavivirus NS1 (28–30)) was diluted 1 in 4 with Da Vinci Green diluent and applied to the sections overnight at room temperature. Sections were washed three times in TBS-T and endogenous peroxidase was blocked by incubating slides in Biocare Medical Peroxidased 1 for 5 minutes. Sections were washed three times in TBS-T and Perkin Elmer Opal HRP Polymer or Perkin Elmer Goat anti-mouse HRP diluted 1:500 in TBST was applied for 60 minutes. Sections were washed three times in TBS-T and signal developed in Vector Nova Red for 5 minutes after which they were washed three times in water. Sections were lightly counterstained in Haematoxylin (program 7 Leica Autostainer), washed in water, dehydrated through ascending graded alcohols, cleared in xylene, and mounted using DePeX or similar.

Apoptosis was detected using the ApopTag Peroxidase In Situ Apoptosis Detection Kit (Merck Catalogue No. S7100) as per manufacturer’s instructions.

For anti-GFAP IHC, antigen retrieval was performed in 0.1M citric acid buffer (pH 6.0) at 1005°C for 20 min. Endogenous peroxidase activity was blocked using 1.0% H_2_O_2_ and 0.1% sodium azide in TBS for 10 min. Endogenous mouse Ig was blocked by incubating sections with goat anti-mouse Fab fragments (Jackson Immunoresearch) diluted 1:50 in Biocare Medical Mouse block M for 60 minutes. Nonspecific antibody binding was inhibited by incubation with Biocare Medical Background Sniper + 2% BSA for 30 min. Mouse anti-GFAP clone GA-5 (Biocare Medical, CM065C), was diluted 1:250 in the above buffer and incubated on sections for 1 hr at room temperature. After washes, Perkin Elmer Goat anti-mouse HRP diluted 1:500 in TBST was applied for 60 minutes. Nova Red development and counter staining was performed as for 4G4 above.

### Infection of human cortical brain organoids

hBOs were reprogrammed from adult dermal fibroblasts (HDFa, Gibco, C0135C) using the CytoTune-iPS 2.0 Sendai Reprogramming Kit (Invitrogen, A16518) (31), and were grown using the CelVivo Clinostar incubator (Invitro Technologies) as described (32). On the day of infection, ∼30-day-old hBOs were transferred from each Clinoreactor into an ultra-low-binding 24 well plate (one hBO per well), and each hBO was infected with 10^5^ CCID_50_ of either JEV_Nakayama_, JEV_FU_, JEV_NSW/22_, MVEV_TC123130_, Imojev (33), or YFV 17D (30) for ∼4 hrs. For shorter term culture (up to 4 days for viral growth kinetics), virus inoculum was removed and hBOs were washed twice with media in the well, before 1 ml differentiation media was added to each well and the 24 well plate was placed within a humidified tissue culture incubator at 37°C, 5% CO_2_ for up to 4 days. For culture up to 11 dpi, hBOs were washed twice with media by transferring them to an ultra-low-binding 6 well plate containing 5 ml of media, and then hBOs were transferred into 50 mm LUMOX gas exchange dishes (SARSTEDT) (4 organoids per dish) containing 7 ml of differentiation media, and placed within a humidified tissue culture incubator at 37°C, 5% CO_2_ for up to 11 days. hBOs were imaged using an EVOS FL (Advanced Microscopy Group), and organoid 2D image circumference was determined by drawing around the edge of the organoid using Image J v1.53 (34).

### Human serum neutralisation assays

Cohort 1 is human serum samples from 9 participants (age >50 years), and were collected ∼28 days (± 2 days) after vaccination with the Imojev vaccine (Sanofi-Aventis Australia). Serum was also collected pre-Imojev vaccination for these same participants for baseline analysis.

Cohort 2 is human serum samples from 10 participants (age range 24 to 60 years), collected at variable times post-vaccination (∼ 2 months to 1 year). Pre-vaccination serum was not collected for Cohort 2. Neutralizing antibodies against Imojev, JEV_Nakayama_, JEV_FU_, JEV_NSW/22_, and MVEV_TC123130_ were measured by plaque reduction neutralization (PRNT) assay (35, 36). BHK-21 cells were seeded at 1.6×10^5^ cells per well in 24 well plates overnight at 37°C. Serum samples were heat-inactivated (56°C, 30 min) and serially diluted 4-fold from 1:5 to 1:160 in BHK-21 cell culture media. Serum was then incubated with 100–110 pfu of Imojev, JEV_Nakayama_, JEV_FU_, JEV_NSW/22_, and MVEV_TC123130_ for 1 h at 37°C. Serum plus virus mixtures were then added to BHK-21 cell monolayers and incubated for 1 hr at 37°C to enable non-neutralized virus to adsorb to cells. Thereafter, 1 ml of 0.375% w/v carboxymethyl cellulose (CMC, Sigma-Aldrich)/RPMI 1640 was added, and the plates were incubated at 37°C in a CO_2_ incubator for 3 days for JEV_Nakayama_ and 4 days for JEV_NSW/22_. The CMC medium was then removed, and the cell monolayers were fixed and stained with 0.1% w/v crystal violet (Sigma-Aldrich) in formaldehyde (1% v/v) and methanol (1% v/v). Plate wells were washed with tap water, dried and the plaques were counted. The PRNT_50_ titer was interpolated from plaque count compared to the average plaque count for the naïve or no serum control.

### Envelope protein structure visualisations

JEV envelope protein structure was downloaded from the Protein Data Bank (PDB: 5WSN) (37). Envelope structure visualisations were generated using PyMol Molecular Graphics System (version 2.3.3; Schrodinger, NY, USA). Virus sequences were downloaded from GenBank (see Supplementary Figure 1 for accession numbers) and aligned using Mega X (Molecular Evolutionary Genetics Analysis 10, Penn State University, State College, PA, USA) and the ClustalW plugin with default parameters. Differences in amino acids compared to Imojev were colored according to structural conservation as described (38).

### Statistics

Statistical analyses of experimental data were performed using IBM SPSS Statistics for Windows, Version 19.0 (IBM Corp., Armonk, NY, USA). The t-test was used when the difference in variances was <4, skewness was > –2 and kurtosis was <2. Otherwise, the non-parametric Kolmogorov-Smirnov test or Mann-Whitney test was used. Paired t-test was used for comparing neutralisation of different viruses with the same human serum sample. For matched human serum samples, a paired non-parametric Wilcoxon matched-pairs signed rank test (GraphPad Prism 8) was used since difference in variance was >4. Kaplan-Meier statistics were determined by log-rank (Mantel-Cox) test. Area under the curve analyses were performed in GraphPad Prism 8, with area under the curve values then compared by t-test. Correlation analyses of PRNT_50_ with envelope conservation used the non-parametric Spearman’s rank-order correlation.

## Results

### JEV_NSW/22_ infection produces a viremia but is not lethal in C57BL/6J mice

The JEV_NSW/22_ G4 virus was isolated from the brain of a stillborn piglet in New South Wales (NSW), Australia in February 2022 (GenBank accession OP904182). The JEV_Nakayama_ genotype 3 prototype was isolated in Japan in 1935 (GenBank accession EF571853), and the JEV_FU_ genotype 2 was isolated in Australia in 1995 (39, 40) (GenBank accession AF217620). MVEV was isolated in Australia in 1974 (41) (MVEV_TC123130_, GenBank accession JN119814). The latter three viruses were isolated from human patients. Brief descriptions of the viral isolates and their phylogenic relationships with other flaviviruses are provided in Supplementary Figure 1.

Six week old adult C57BL/6J mice were infected subcutaneously (s.c.) with 5×10^5^ CCID_50_ of the aforementioned viruses. Viremia for all four viruses were broadly similar, peaking 1 day post infection (dpi) at 2-3 log_10_CCID_50_/ml of serum, with nearly all mice showing no detectable viremia by day 4 (Figure 1A-D). Infection appeared to stall weight gain for most mice until day ∼10 (Figure 1E). Between 8 to 12 dpi, four mice (2 infected with JEV_Nakayama_, 1 JEV_FU_ and 1 MVEV_TC123130_) out of the total of 24 mice showed weight loss ≥20% and were euthanized (Figure 1E, †). A further four mice (1 JEV_Nakayama_, 2 JEV_FU_, 1 MVEV_TC123130_) lost >5% of their body weight, but subsequently recovered. None of the JEV_NSW/22_ infected mice lost more than ∼3% of their body weight (Figure 1E). Kaplan-Meier survival curves provided no significant differences for the different viral isolates (Figure 1F). Mice that were euthanized also displayed varying levels of abnormal posture (hunching), reduced activity, and fur ruffling, on the day of euthanasia (Supplementary Figure 2A). For the 4 mice that were euthanized (Figure 1E), these mice had very high levels of brain infection (≈8-9 log_10_CCID_50_/mg) (Figure 1G). At the time of euthanasia, viral titers in the spleen in these mice were below the level of detection (Figure 1G), consistent with the viremia data.

**Figure 1.**
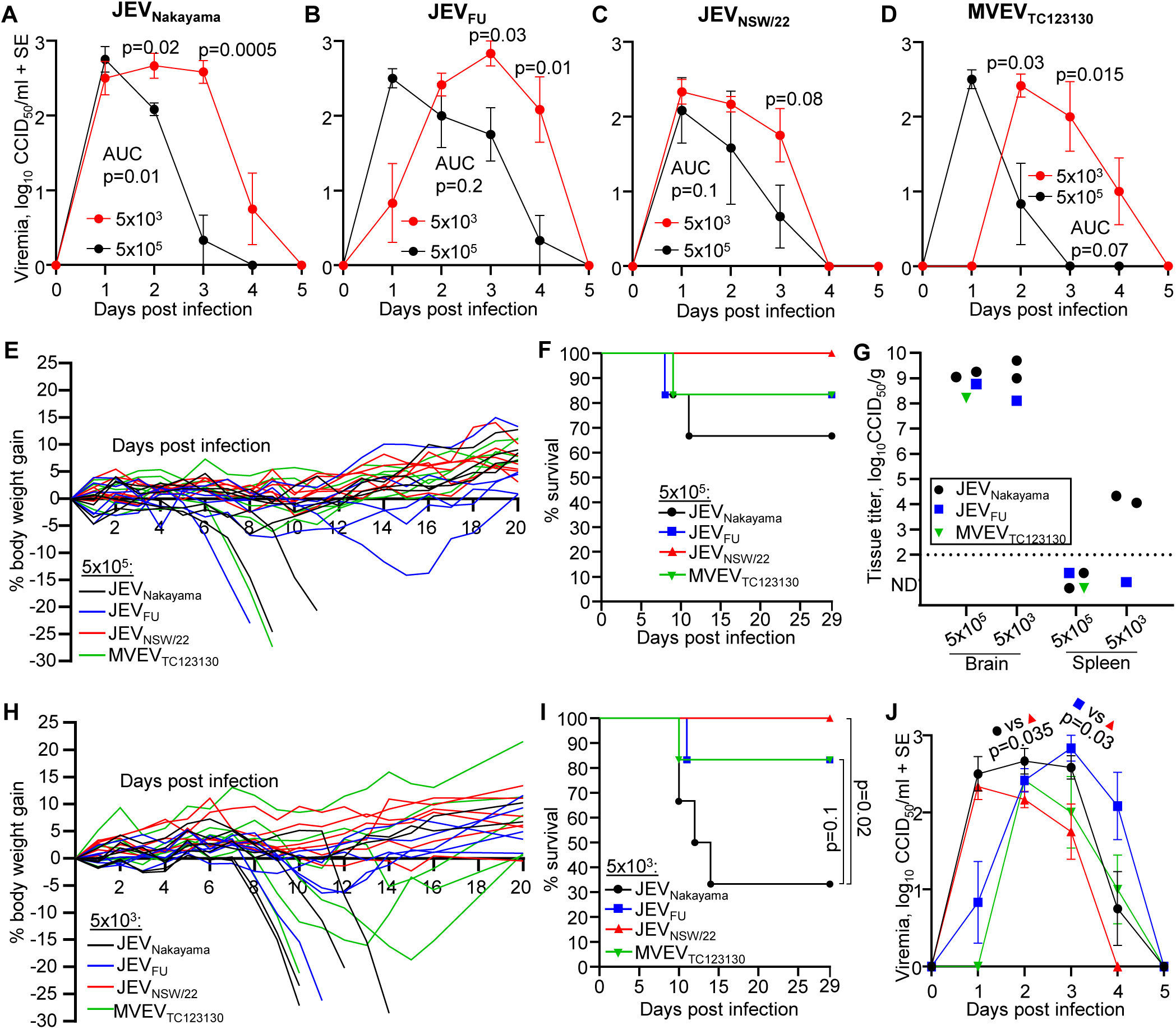
JEV and MVEV infection in C57BL/6J mice. (A-D) Female ≈6 week old C57BL/6J mice were infected s.c. with 5×10^5^ (black lines) or 5×10^3^ (red lines) CCID_50_ of the indicated viruses. Mean viremia for n=6 per group over 5 days is shown. Error bars represent standard error. All mice recorded a detectable viremia for at least one timepoint. Limit of detection is 2 log_10_CCID_50_/ml of serum. Statistics represent t-test or Kolmogorov-Smirnov test at the indicated timepoint or for area under the curve (AUC) values (see methods section). (E) Percent body weight change of individual mice after infection with the indicated virus at 5×10^5^ CCID_50_ compared to each mouse’s weight on day zero. Four mice lost ≥20% body weight and were euthanized (†). (F) Kaplan-Meier plot showing percent survival (n=6 for each virus/virus isolate inoculated at 5×10^5^ CCID_50_). (G) Viral tissue titers in brains and spleens of seven euthanized mice at the time when the criteria for humane euthanasia was met (see Supplementary Figure 2) (n=4 JEV_Nakayama_ – black circles, n=2 JEV_FU_ – blue square, n=1 MVEV_TC123130_ green downward triangle). Tissue titers determined by CCID_50_ assay (limit of detection ∼2 log_10_CCID_50_/g). (H) Percent body weight change of individual mice after infection with the indicated virus at 5×10^3^ CCID_50_ compared to each mouse’s weight on day zero. Six mice lost ≥20% body weight and were euthanized (†). (I) Kaplan-Meier plot showing percent survival (n=6 for each virus/virus isolate inoculated at 5×10^3^ CCID_50_). (J) Viremia comparing JEV_Nakayama_ (black circles), JEV_FU_ (blue squares), JEV_NSW/22_ (red triangles) and MVEV_TC123130_ (green downward triangles) at 5×10^3^ inoculation dose; data is a reanalysis of data presented in Figure 1A-D. Data is mean of n=6 per group and error bars represent standard error. Statistics are t-test for JEV_Nakayama_ versus JEV_NSW/22_ on day 2 and JEV_FU_ versus JEV_NSW/22_ on day 3.

We then infected six week old adult C57BL/6J mice with 5×10^3^ CCID_50_ (s.c.) of the aforementioned viruses to determine if reduced inoculation dose can increase viremia and neuropenetrance, as has been reported previously (42). Compared to mice infected with 5×10^5^ CCID_50_, viremia in mice infected with 5×10^3^ CCID_50_ of virus was significantly higher at later time points for JEV_Nakayama_ (Figure 1A, day 2-3), JEV_FU_ (Figure 1B, day 3-4), and MVEV_TC123130_ (Figure 1D, day 2-3). For JEV_NSW/22_, viremia at day 3 post infection was higher on average but this did not reach statistical significance (Figure 1C). The viremia area under the curve was significantly higher for JEV_Nakayama_ inoculated with the lower dose (Figure 1A), but was not significantly different for the other viruses. This coincided with an increase in mortality for JEV_Nakayama_ inoculated at the lower dose, but not for the other viruses (Figure 1H-I). Viral titers in the brains or spleens of mice that succumbed to infection were not significantly different between inoculum doses (Figure 1G). JEV_NSW/22_ had a significantly lower viremia compared to JEV_Nakayama_ and JEV_FU_ (Figure 1J), consistent with significantly lower mortality compared with JEV_Nakayama_ (Figure 1I), with no C57BL/6J mice infected with JEV_NSW/22_ succumbing to infection (Figure 1F and I). Overall, this suggests that JEV_NSW/22_ was significantly less virulent in C57BL/6J mice.

### C57BL/6J mice infected with JEV or MVEV led to viral neuroinvasion, apoptosis and reactive astrogliosis

The brains of C57BL/6J mice were analysed by immunohistochemistry (IHC) using the pan-flavivirus anti-NS1 antibody (4G4). This antibody detects the viral non-structural protein 1, which is a highly conserved multi-functional protein important for flavivirus replication (43). Staining was consistently seen in the cortex for JEV and MVEV infected mice, with staining also seen in the thalamus (Figure 2A, Supplementary Figure 3A-C). Prevailing infection of the cerebral cortex and thalamus parallels IHC data from post-mortem human brains (6, 44). In some mice, virus was detected in other brain regions, including the hippocampus, anterior olfactory nucleus (Figure 2A, JEV_Nakayama_), and caudate putamen (Supplementary Figure 3B, JEV_FU_). A JEV_FU_ infected mouse that lost ∼15% body weight and then recovered (Figure 1E) had residual virus staining in the cortex at day 29 post infection (Supplementary Figure 3D), while a mouse that survived infection with minimal weight loss had no detectable viral antigen staining (Supplementary Figure 3E). As expected, 4G4 staining was associated primarily with cells showing neuronal morphology (Figure 2A, Supplementary Figure 3D).

**Figure 2.**
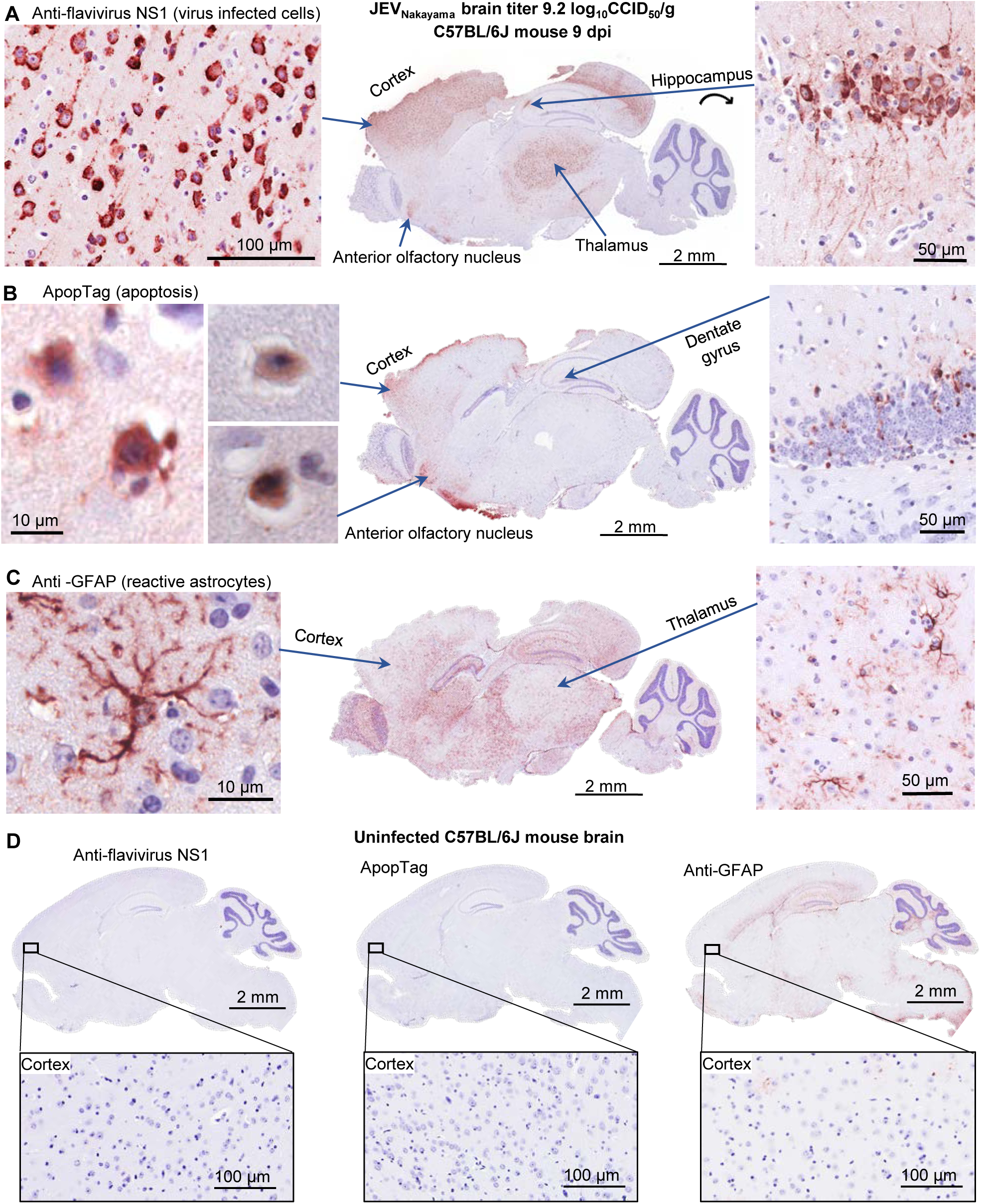
IHC for viral antigen, apoptosis and reactive astrocytes in JEV infected C57BL/6J mouse brain. IHC of JEV_Nakayama_ infected brain (required euthanasia on 9 dpi, brain virus titer 9.2 log_10_CCID_50_/g), which is representative of all other C57BL/6J brains from infected mice requiring euthanasia (n=10). (A) Staining for flavivirus NS1 using the 4G4 monoclonal antibody. High magnification images show cells with neuronal morphology in the cortex (left) and hippocampus (right). The latter also shows staining of dendrites and axons (fibrilar patterns above and densely staining cells). 4G4 staining of brains of the other mice requiring euthanasia (marked by † in Figure 1E) is shown in Supplementary Figure 3. (B) ApopTag staining of the same brain as in ‘A’. High magnification images show cells with neuronal morphology in the cortex (left) and apoptotic cells in the dentate gyrus (right). ApopTag staining of the brains of the other mice requiring euthanasia is shown in Supplementary Figure 4. (C) Staining for GFAP, a marker of reactive astrocytes, for the same brain as in ‘A’. High magnification images show a typical reactive astrocyte in the cortex (left) and reactive astrocytes in the thalamus (right). GFAP IHC for the brains of the other mice requiring euthanasia is shown in Supplementary Figure 5. (D) IHC negative controls; uninfected mouse brain stained with 4G4 (left), ApopTag (middle), and anti-GFAP (right).

ApopTag staining was evident in areas that were positive for virus staining, particularly in the cortex, and was again associated with cells showing neuronal morphology (Figure 2B and Supplementary Figure 4A-C). Apoptotic cells were also evident in the mouse that showed ∼15% body weight loss and recovered (Supplementary Figure 4D). Mice with signs of disease from JEV or MVEV infection (Supplementary Figure 2) thus showed high levels of viral antigen and apoptosis in the brain. In addition, like in humans, animals can recover from brain infection, despite apoptotic damage, although neurological sequelae may ensue.

Astrocytes are a type of glial cell that provides physical and chemical support to neurons. Astrocytes become activated (reactive astrogliosis) in response to brain pathology, including from JEV infection (45–47). Reactive astrocytes are characterized by the upregulated expression of glial fibrillary acidic protein (GFAP) and the growth/extension of cellular processes (48).

Brains from mice with signs of disease from JEV or MVEV infection (Supplementary Figure 2) had significant upregulation of GFAP positive astrocytes throughout the brain, including the heavily infected areas of the cortex, thalamus and anterior olfactory nucleus (Figure 2C, Supplementary Figure 5). GFAP is constitutively expressed in astrocytes of the hippocampus, corpus callosum, and cerebral peduncle (49), consistent with staining seen in uninfected brains (Figure 2D, right). Widespread reactive astrogliosis is thus a feature of lethal JEV and MVEV encephalitis in C57BL/6J mice.

### JEV_NSW/22_ is less virulent than JEV_Nakayama_ and JEV_FU_ in Irf7^-/-^ mice

C57BL/6J mice had a relatively low viremia and low penetrance of lethal brain infection for JEV_NSW/22_ (Figure 1). A model with a higher penetrance of brain infection is clearly desirable, given encephalitis is the key clinical concern for JEV infections. Interferon regulatory factor 7 knockout (*Irf7^-/-^* mice) were thus chosen as these mice enhanced neuroinvasion for WNV (50), as well as another arbovirus (24). We first sought to optimize the inoculation dose for JEV_NSW/22_ in *Irf7^-/-^* mice by comparing 5×10^5^, 5×10^4^, 5×10^3^, and 5×10^2^ CCID_50_. The inoculation dose of 5×10^3^ CCID_50_ provided the highest viremia area under the curve, which was significantly higher than 5×10^4^ and 5×10^5^ CCID_50_ inoculation doses (Figure 3A). The highest infection dose (5×10^5^) led to the highest peak viremia on day 1 post infection, but was cleared significantly more quickly than lower infection doses (Figure 3A, p=0.004). Our data agrees with the concept of an ‘optimal dose’ that is not too high as to excessively stimulate type-I IFN, but is high enough to establish a robust viremia (42). One mouse infected with 5×10^3^ CCID_50_ JEV_NSW/22_ succumbed to infection (Figure 3C-D), while all other mice survived infection. An inoculation dose of 5×10^3^ CCID_50_ was thus chosen to compare JEV and MVEV strains in *Irf7^-/-^* mice.

**Figure 3.**
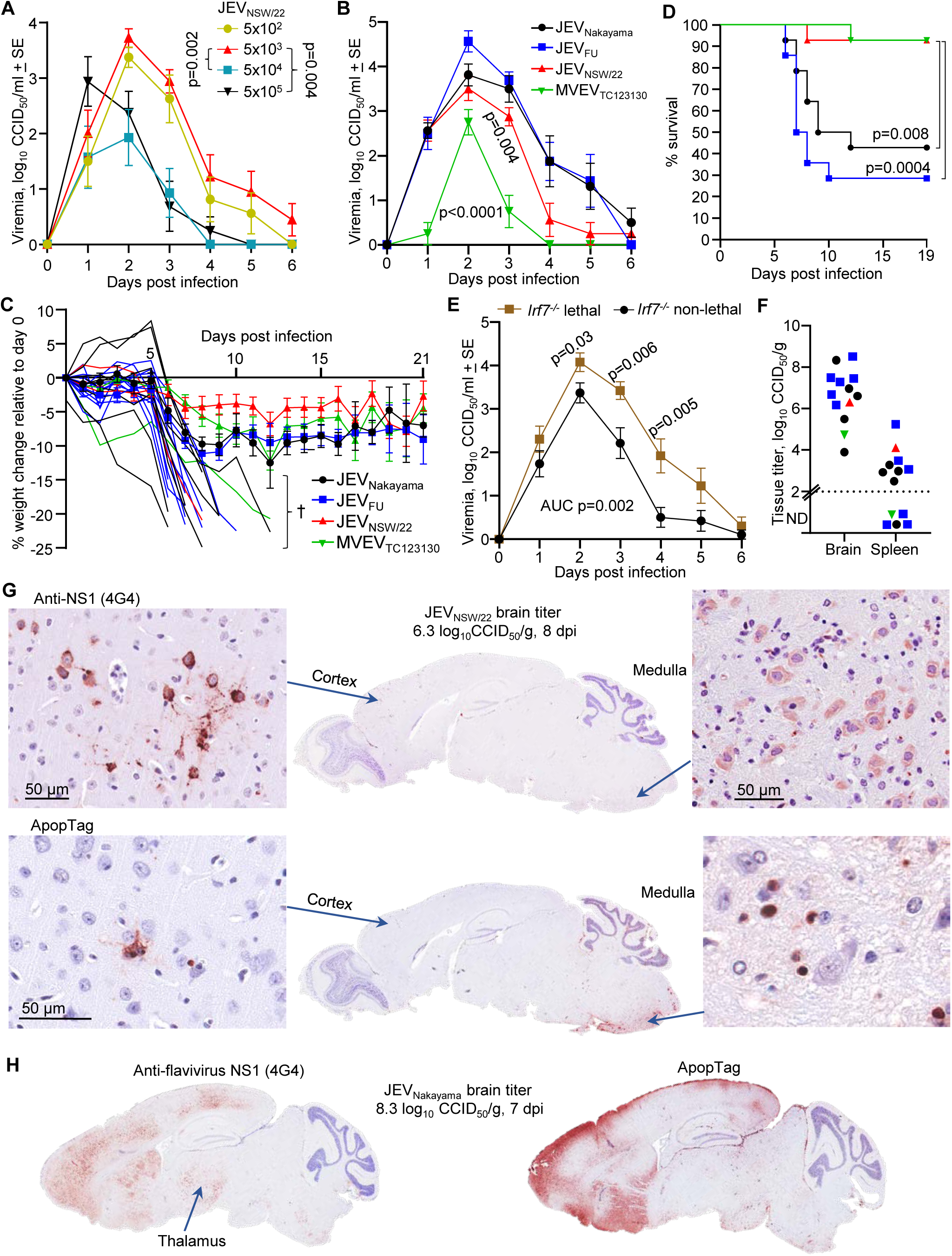
JEV neuroinvasive infection in *Irf7^-/-^* mice. (A) *Irf7^-/-^* mice (15-48 weeks old) were infected s.c. with 5×10^5^, 5×10^4^, 5×10^3^ or 5×10^2^ CCID_50_ of JEV_NSW/22_. Data is the mean of n=6 per group and error bars represent standard error. Statistics are a t-test of area under the curve for the indicated comparisons. (B) *Irf7^-/-^* mice (15-48 weeks old) were infected s.c. with 5×10^3^ of the indicated virus (n=8 for each virus). Statistics are a t-test of area under the curve for JEV_FU_ versus JEV_NSW/22_ or MVEV_TC123130_. (C) Percent body weight change compared to 0 dpi. 20 mice lost >20% body weight or reached a disease score that required euthanized (marked by †) and are plotted individually. The weight change for the remaining mice are shown as means ± SE. Data is from three independent experiments, total n=14 mice per group. (D) Kaplan-Meier plot showing percent survival (n=14 per group, data from three independent experiments). Statistics by log-rank (Mantel-Cox) tests. Symbols as for C. (E) Viremia of *Irf7^-/-^* mice averaged for mice with non-lethal outcomes (black circles, n=19), versus those with lethal outcomes (brown squares, n=13). Statistics are comparing average viremia of mice with lethal outcomes versus mice with non-lethal outcomes at each timepoint by t-test or Kolmogorov-Smirnov test. T-test of area under the curve values for this comparison is also shown. (F) Brain and spleen tissue titers for 13 euthanized mice at the time when the criteria for humane euthanasia was met (see Supplementary Figure 6) (n=5 JEV_Nakayama_, n=6 JEV_FU_, n=1 JEV_NSW/22_, n=1 MVEV_TC123130_). Tissue titers determined by CCID_50_ assay (limit of detection ∼2 log_10_CCID_50_/g). (G) IHC of JEV_NSW/22_ infected brain (euthanized on day 8, brain virus titer 6.3 log_10_CCID_50_/g) using 4G4 monoclonal antibody (top) or ApopTag (bottom). High magnification images of cortex (left) and medulla (right). (H) IHC using 4G4 (left) and ApopTag IHC (right) for the JEV_Nakayama_ infected brain with the highest virus titer (euthanized on day 7, brain virus titer 8.3 log_10_CCID_50_/g). IHC images are representative of n=20 brains of mice that succumbed to infection (some other examples shown in Supplementary Figure 8).

JEV_NSW/22_, JEV_Nakayama_ and JEV_FU_ infection of *Irf7^-/-^* mice (dose 5×10^3^ CCID_50_) resulted in robust viremias, while MVEV_TC123130_ viremia was significantly lower than JEV (Figure 3B, p<0.0001). JEV_NSW/22_ had a significantly lower viremia (area under the curve) compared to JEV_FU_ (Figure 3E, p=0.004). Infection of *Irf7^-/-^* mice with the different JEV isolates and MVEV (n=14 mice per group, n=56 total infected mice) illustrated a mean weight loss of 5-10% for surviving mice across all four groups (Figure 3C), with the exception of 20 mice that reached ethically defined end points for weight loss and/or disease manifestations (Supplementary Figure 6). Kaplan-Meier curves illustrated significantly higher survival for *Irf7^-/-^* mice infected with JEV_NSW/22_ or MVEV_TC123130_ when compared with JEV_Nakayama_ and JEV_FU_, with only 1/14 JEV_NSW/22_ or MVEV_TC123130_ infected *Irf7^-/-^* mice requiring euthanasia (Figure 3D). These data are consistent with the significantly lower viremias of JEV_NSW/22_ and MVEV_TC123130_ infected *Irf7^-/-^*mice (Figure 3B). On average, mice that succumbed to infection had a more robust viremia (Figure. 3E), further implicating a higher viremia in a higher chance of lethal neuropenetrance in *Irf7^-/-^*mice.

Brain titers in euthanized *Irf7^-/-^* mice were ∼4-9 log_10_CCID_50_/gram, while spleen titers were <2-5 log_10_CCID_50_/g (Figure 3F). IHC staining for viral antigen confirmed JEV infection in the brains of *Irf7^-/-^* mice, with reduced levels of staining reflecting lower brain titers (Figure 3G-H, Supplementary Figure 7). Viral antigen staining was consistently present in the cortex, with staining also seen in the medulla, thalamus, and hippocampus (Figure 3G-H, Supplementary Figure 7). Brain regions showing ApopTag staining broadly overlapped with staining for viral antigen (Figure 3G-H). Reactive astrogliosis was also evident (Supplementary Figure 5).

Overall, this suggests that JEV infection of *Irf7^-/-^* mice produced a consistent viremia, with mice showing brain infection and apoptotic damage.

### JEV_NSW/22_ and JEV_FU_ are more sensitive to type I IFN compared to JEV_Nakayama_ and MVEV_TC123130_

Viremia and survival were compared between C57BL/6J and *Irf7^-/-^* mouse strains infected with 5×10^3^ CCID_50_ virus to determine whether differences in viremia may be associated with the differences in viral neuropenetrance and survival. Area under the curve analyses showed a significantly increased/prolonged viremia in *Irf7^-/-^* mice compared to C57BL/6J mice for all JEV strains (Figure 4A-C). However, MVEV_TC123130_ viremia was not significantly improved in *Irf7^-/-^* mice (Figure 4D). Kaplan-Meier survival curves indicated that JEV_FU_ had a significantly increased mortality in *Irf7^-/-^* mice compared to C57BL/6J mice (Figure 4F), while there was no significant difference between mouse strains for the other viruses (Figure 4E, G-H).

**Figure 4.**
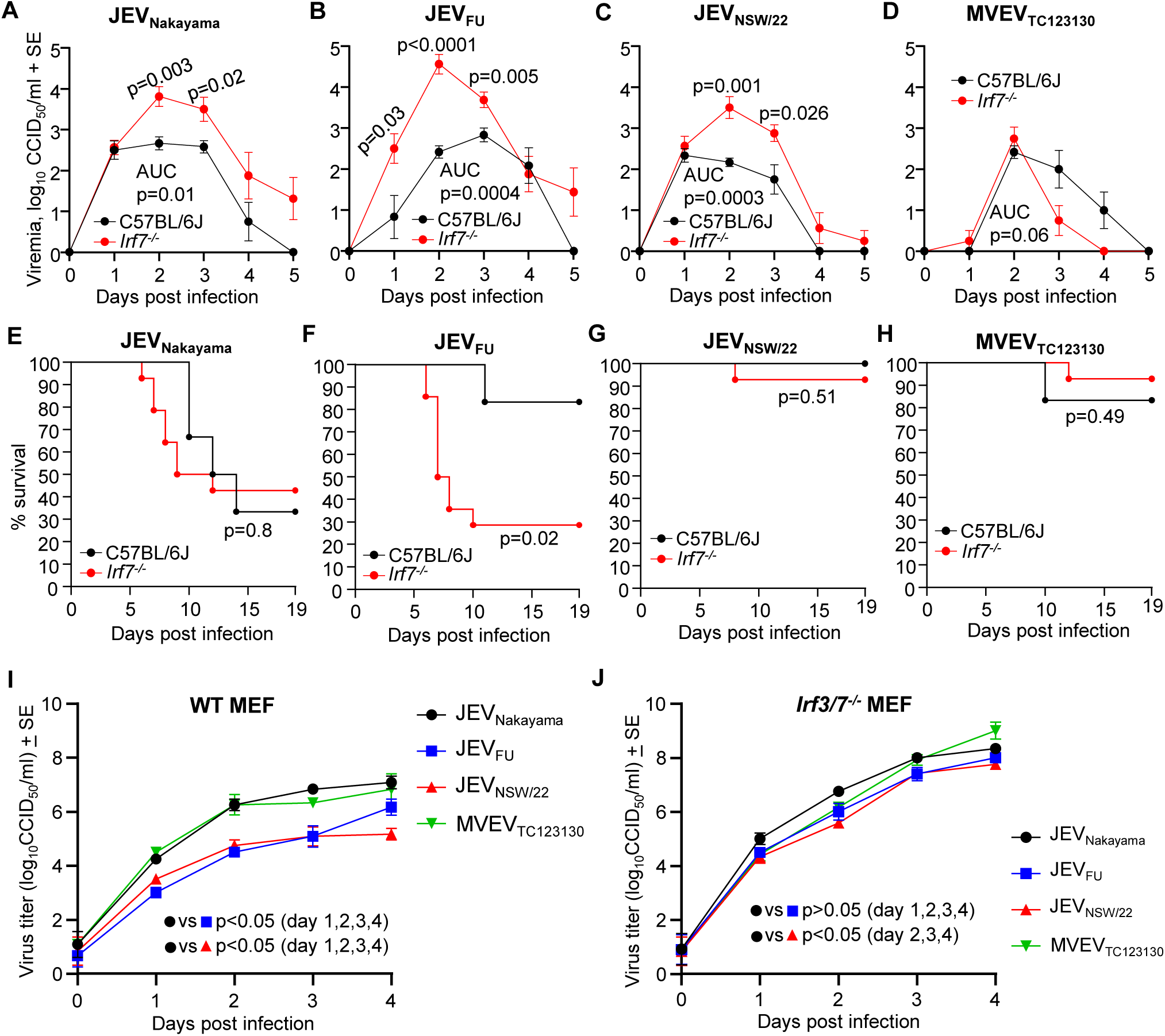
Comparisons of viremia and survival between C57BL/6J versus *Irf7^-/-^* mice, and comparison of virus replication in wild-type versus *Irf3/7^-/-^* mouse embryonic fibroblasts (MEFs). C57BL/6J or *Irf7^-/-^*mice were infected with 5×10^3^ CCID_50_ of JEV_Nakayama_, JEV_FU_, JEV_NSW/22_, or MVEV_TC123130_. Data shown in panels A-H are a re-analysis of data presented in Figure 1A-D, Figure 1I, Figure 3B, and Figure 3D. (A-D) Female ≈6 week old C57BL/6J mice (black lines) or *Irf7^-/-^* mice (red lines) were infected s.c. with 5×10^3^ CCID_50_ of the indicated viruses. Mean viremia for n=6 C57BL/6J mice and n=8 Irf7^-/-^ mice per group over 5 days is shown. Error bars represent standard error. Limit of detection is 2 log_10_CCID_50_/ml of serum. Statistics represent t-test or Kolmogorov-Smirnov test at the indicated timepoint or for area under the curve (AUC) values (see statistics methods section). (E-H) Kaplan-Meier plot showing percent survival for C57BL/6J mice (black line, n=6) and Irf7^-/-^ mice (ref line, n=14) infected with 5×10^3^ CCID_50_ of the indicated virus. Statistics by log-rank (Mantel-Cox) tests. (I) Wild type or (J) *Irf3/7^-/-^*MEFs were infected with JEV_Nakayama_ (black circles), JEV_FU_ (blue squares), JEV_NSW/22_ (red triangles), or MVEV_TC123130_ (green downward triangles) at MOI 0.1. Virus titer in the culture supernatant was monitored over 4 days. Data is the mean two independent experiments with a total of n=6 replicates per group. Error bars represent standard error. Statistics are by t-test or Kolmogorov-Smirnov test for the indicated comparisons.

Differences in sensitivity to type I IFN between virus strains may explain the different outcomes in C57BL/6J versus *Irf7^-/-^* mice. To examine type I IFN sensitivity in the absence of *in vivo* adaptive immune responses, mouse embryonic fibroblasts (MEFs) with (wild-type) and without (*Irf3/7^-/-^*) functional type I IFN production were infected with JEV_Nakayama_, JEV_FU_, JEV_NSW/22_ and MVEV_TC123130_. In wild-type MEFs, JEV_FU_ and JEV_NSW/22_ had significantly lower virus replication kinetics compared to JEV_Nakayama_ and MVEV_TC123130_ (Figure 4I). In *Irf3/7^-/-^* MEFs, the difference in replication between virus strains was diminished, with JEV_FU_ not significantly different to JEV_Nakayama_ (Figure 4J). This suggests that JEV_FU_ and JEV_NSW/22_ were more sensitive to type I IFN compared to JEV_Nakayama_ and MVEV_TC123130_, consistent with an increase in mortality in *Irf7^-/-^* mice (partially defective type I IFN responses) compared to C57BL/6J mice for JEV_FU_ and JEV_NSW/22_ but not for JEV_Nakayama_ or MVEV_TC123130_.

### Ifnar^-/-^ mice are a model of lethal JEV viremia without neuroinvasion

As JEV_NSW/22_ did not lead to fatal neuroinvasion in C57BL/6J mice, and fatal neuroinvasion in Irf7^-/-^ mice was rare, type I interferon receptor knockout (*Ifnar^-/-^*) mice were evaluated. *Ifnar^-/-^* mice have been used extensively in studies of pathogenic flaviviruses such as Zika virus (ZIKV), West Nile virus (WNV), and Yellow fever virus (YFV), and generally provide a robust viremia and a lethal outcome (21, 29, 30, 51). Adult *Ifnar^-/-^* mice were infected with 5×10^5^ CCID_50_ of JEV_Nakayama_, JEV_FU_, JEV_NSW/22_, or MVEV_TC123130_ via s.c. injection. A robust viremia developed, reaching 6-8 log_10_CCID_50_/ml of serum by 2 dpi (Figure 5A). JEV_NSW/22_ infected mice had ≈2 log lower viremia compared to JEV_Nakayama_ and JEV_FU_ on 3 dpi (Figure 5A). Mice displayed a rapid loss of body weight, which was slightly delayed for JEV_NSW/22_ and MVEV_TC123130_ at 2 dpi (Figure 5B), requiring euthanasia 2/3 dpi (Figure 5C). Mice infected with any of the JEV strains also displayed varying levels of abnormal posture (hunching), reduced activity, and fur ruffling (Supplementary Figure 8A).

**Figure 5.**
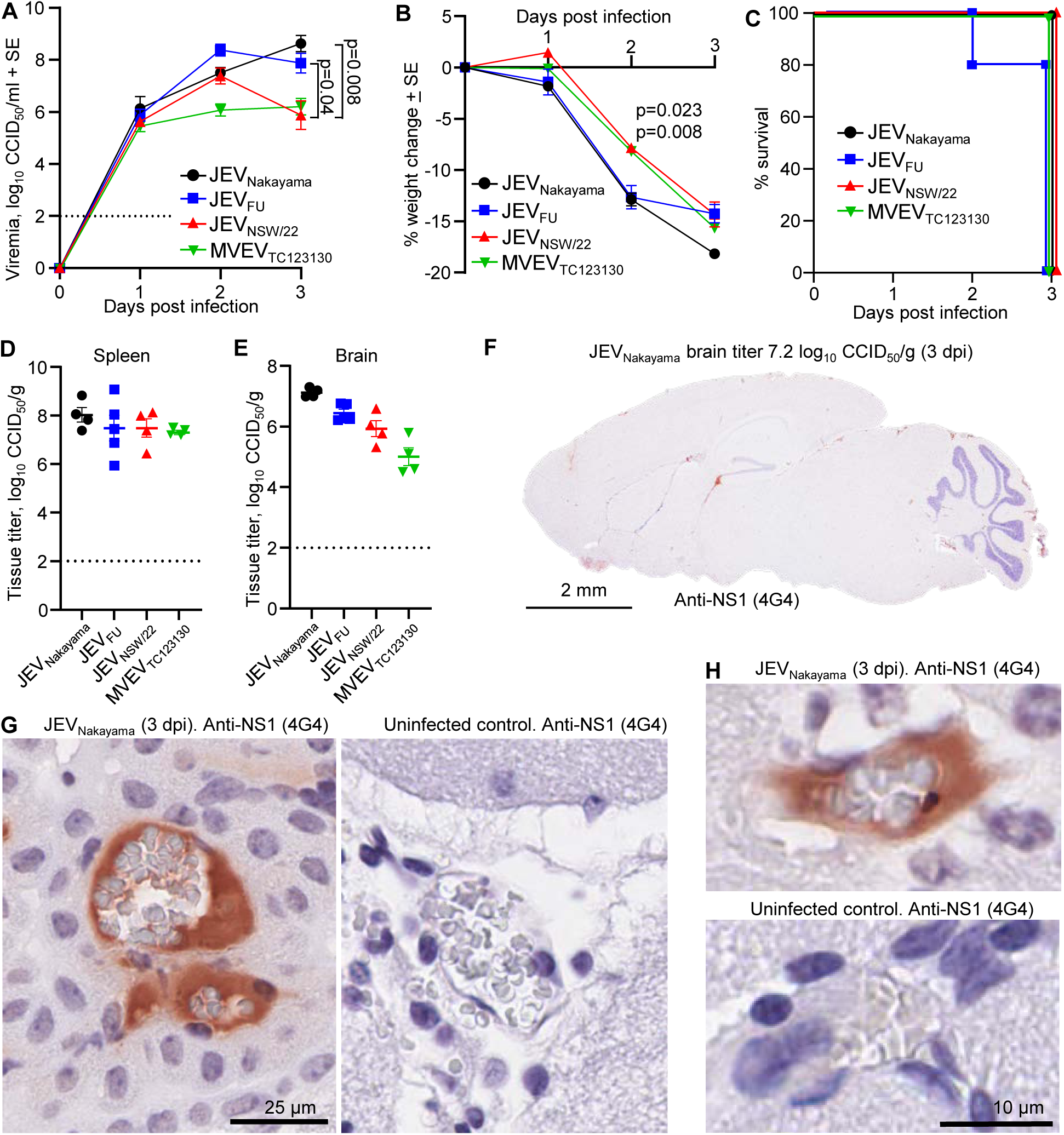
JEV and MVEV lethal viremia in *Ifnar^-/-^* mice. Female *Ifnar^-/-^* mice (9-24 week old) were infected s.c. with 5×10^5^ CCID_50_ of the indicated virus (n=4 for JEV_Nakayama_, JEV_NSW/22_ and MVEV_TC123130_, and n=5 for JEV_FU_). (A) Mean viremia determined by CCID_50_ assay (limit of detection 2 log_10_CCID_50_/ml). Statistics by t-tests. (B) Mean percent body weight change compared to 0 dpi. Statistics 2 dpi JEV_NSW/22_ versus JEV_Nakayama_ (Kolmogorov-Smirnov test, p=0.023) and for JEV_NSW/22_ versus JEV_FU_ (t-test, p=0.008). (C) Kaplan-Meier plot showing percent survival. (D) Viral tissue titers in spleens harvested at euthanasia (3 dpi for all mice except for 1 JEV_FU_ mouse at 2 dpi), determined by CCID_50_ assay (limit of detection ∼2 log_10_ CCID_50_/g). (E) Viral titers in brains harvested at euthanasia. (F) IHC staining for flavivirus NS1 using the 4G4 monoclonal antibody. The brain shown was infected with JEV_Nakayama_ (titer 7.2 log10 CCID50/g, 3 dpi). Staining was representative of all other JEV brains. (G, H) High magnification images from F showing NS1 staining in blood vessels surrounding the pale grey, biconcave shaped, red blood cells.

Consistent with the robust viremia (Figure 5A), spleen tissue titers reached ∼6-9 log_10_CCID_50_/g at 2-3 dpi (Figure 5D). Similar levels were seen in brains (Figure 5E); however, 4G4 staining illustrated that brain cells were not infected (Figure 5F). Viremia and brain titers correlated strongly (Supplementary Figure 8B), arguing that the brain titers (Figure 5E) likely arose from virus in the blood vessels of the brain. Interestingly, anti-flavivirus NS1 antibody staining was clearly present in blood vessels (Figure 5G,H), with extracellular JEV NS1 also found in serum of JEV patients (52). Secreted NS1 from JEV infected cells is reported to increase vascular leakage and may contribute to mortality (53), although overt vascular leakage of NS1 was not evident in IHC, with lack of type I IFN signalling potentially involved (54, 55). Neither apoptosis (Supplementary Figure 8C), nor significantly increased staining for reactive astrocytes (Supplementary Figure 5, *Ifnar^-/-^*) were apparent for brains of JEV-infected *Ifnar^-/-^*mice. Therefore, *Ifnar^-/-^* mice infected with JEV or MVEV represent a model of lethal viremia without viral neuroinvasion.

### JEV causes severe histological lesions in C57BL/6J and Irf7^-/-^ mouse brains

H&E staining of JEV infected mouse brains revealed a series of histopathological lesions, predominantly neuronal degeneration with pyknotic nuclei (often associated with apoptosis), neuronal vacuolation, perivascular cuffing, leukocyte infiltrates, hemorrhage, meningitis and microgliosis (Figure 6A). IHC for the microglial marker, Iba1, confirmed the presence of microgliosis (Figure 6B).

**Figure 6.**
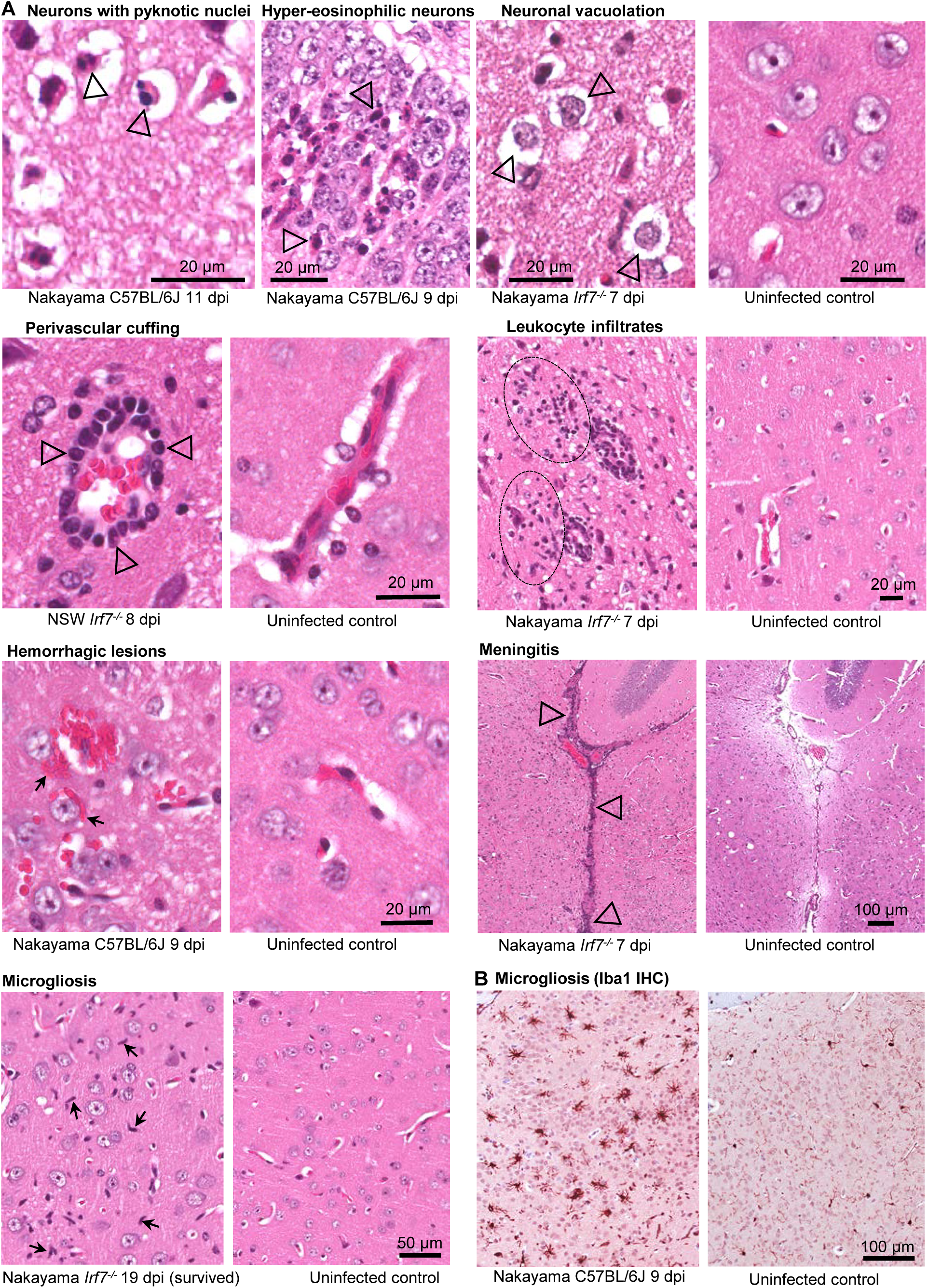
Histological lesions in brains of JEV infected mice. (A) Examples of indicated lesions stained with H&E. Degeneration of neurons indicated by (i) pyknotic nuclei (black unfilled arrow heads indicating condensation and fragmentation of nuclei, staining dark blue), and (ii) hyper-eosinophilic cytoplasm of degenerating neurons in the cornus ammonis of the hippocampus (area also staining for viral antigen Supplementary Figure 10). Neuronal vacuolation indicated by fluid accumulation around the neurons (arrowheads). Perivascular cuffing is indicated by leukocytes aggregating in blood vessels (arrowheads). Leukocyte infiltrates (extravascular) are indicated by dashed ovals. Hemorrhagic lesions are indicated by extravascular red blood cells (arrows). Microgliosis is indicated by accumulation of microglia, which have an elongated rod-shaped nuclei (arrows). Meningitis is indicated by accumulation of leukocytes around the meninges (arrowheads). Images of uninfected controls accompany each image(s) of lesions. Histology scores for all mouse brains are shown in Supplementary Figure 9 and 10. (B) IHC using anti-Iba1, a microglial marker.

The presence of H&E lesions were scored for the different JEV isolates and mouse strains (Supplementary Figure 9). C57BL/6J and *Irf7^-/-^* mice that were euthanized at the time when viral neuroinvasion led to the criteria for humane euthanasia being met (full score cards provided in Supplementary Figure 2 and 7) had a broadly similar incidence of the aforementioned lesions, whereas only hemorrhage was seen in *Ifnar^-/-^* mice (Supplementary Figure 9, Acute). Leukocyte infiltration was quantified by calculating the ratio of nuclear/cytoplasmic staining, as leukocytes have a higher ratio of nuclear to cytoplasmic staining (27). This confirmed that C57BL/6J and *Irf7^-/-^*mice that succumbed to infection had significantly increased leukocyte infiltrates (Supplementary Figure 9B). For C57BL/6J and *Irf7^-/-^* mice that survived and were euthanized later (nominally chronic phase), only hemorrhage was consistently observed (Supplementary Figure 9, Chronic).

Lesions in C57BL/6J and *Irf7^-/-^* mouse brains were associated with areas of high virus infection, most prominently in the cortex (Supplementary Figure 10); however, lesions were also found in other areas of the brain including where there was minimal viral antigen staining (Supplementary Figure 11A). Some mice that lost >8% body weight and then recovered had persistent lesions detectable at the latest time point sampled (up to day 32 for C57BL/6J and day 21 for *Irf7^-/-^*) (Supplementary Figure 11B), consistent with persistent neurological sequelae in humans that survive infection. Mice that did not lose any body weight did not show any overt brain lesions at the later time points, indicating that brain infection causing weight loss is associated with persistent lesions.

Overall, the lesions in C57BL/6J and *Irf7^-/-^* mouse brains are consistent with H&E detectable lesions in post-mortem human JEV infected brains (6, 45, 56–58). To provide additional potential insights into the nature of the cellular infiltrates (e.g. perivascular cuffing and leukocyte infiltrates, Figure 6), immune cell type abundance estimates were obtained from reanalyzed (59) publically available JEV-infected mouse brain RNA-Seq expression data (60) using SpatialDecon (61). The dominant cell types identified were CD4 T cells, NKT cells, monocytes/macrophages, neutrophils, innate lymphoid cells, T regs, and CD8 T cells, with an increase in microglia cells also identified (Supplementary Figure 12). GFAP was significantly up-regulated 5.4 fold by infection (Supplementary Table 3, log_2_FC 2.4), consistent with the GFAP IHC (Figure 2C) and reactive astrogliosis seen herein. Iba1 (Aif1) was also significantly up-regulated 5.5 fold (Supplementary Table 3, log_2_FC 2.5), consistent with the H&E and IHC (Figure 6) and microgliosis observed in our study.

### Productive replication of JEV in human cortical brain organoids

Human cortical brain organoids (hBOs) have been used to investigate the pathogenesis of flaviviruses, including JEV and ZIKV (21, 62, 63). We generated ∼30 day old hBOs from adult human dermal fibroblast (HDFa) cells, growing the hBOs in a CelVivo Clinostar incubator as described (32) (Figure 7A). hBOs were infected with 10^5^ CCID_50_ of the indicated JEV and MVEV isolates, as well as (i) the Imojev chimeric virus vaccine (previously called ChimeriVax-JE) comprising the prME genes of the attenuated JEV_SA14-14-2_ strain on the YFV 17D backbone (33, 64) and (ii) the Yellow Fever live attenuated vaccine strain (YFV 17D) (30), with wild-type YFV infection (65), and occasionally YFV 17D vaccination (66), able to cause neuropathology. hBOs were fixed in formalin 4 dpi and IHC undertaken using the anti-NS1 monoclonal antibody 4G4 (30). JEV_Nakayama_ infected hBOs showed the most pronounced viral antigen staining, with JEV_NSW/22_, JEV_FU_ and MVEV_TC123130_ also showing clear staining (Figure 7B). Viral antigen was primarily localized to the outer surface of the organoids (Figure 7B), where cells are in direct contact with the culture medium. Viral antigen staining for Imojev and YFV 17D infected hBOs was also seen, but was sparse (Figure 7B and Supplementary Figure 13A).

**Figure 7.**
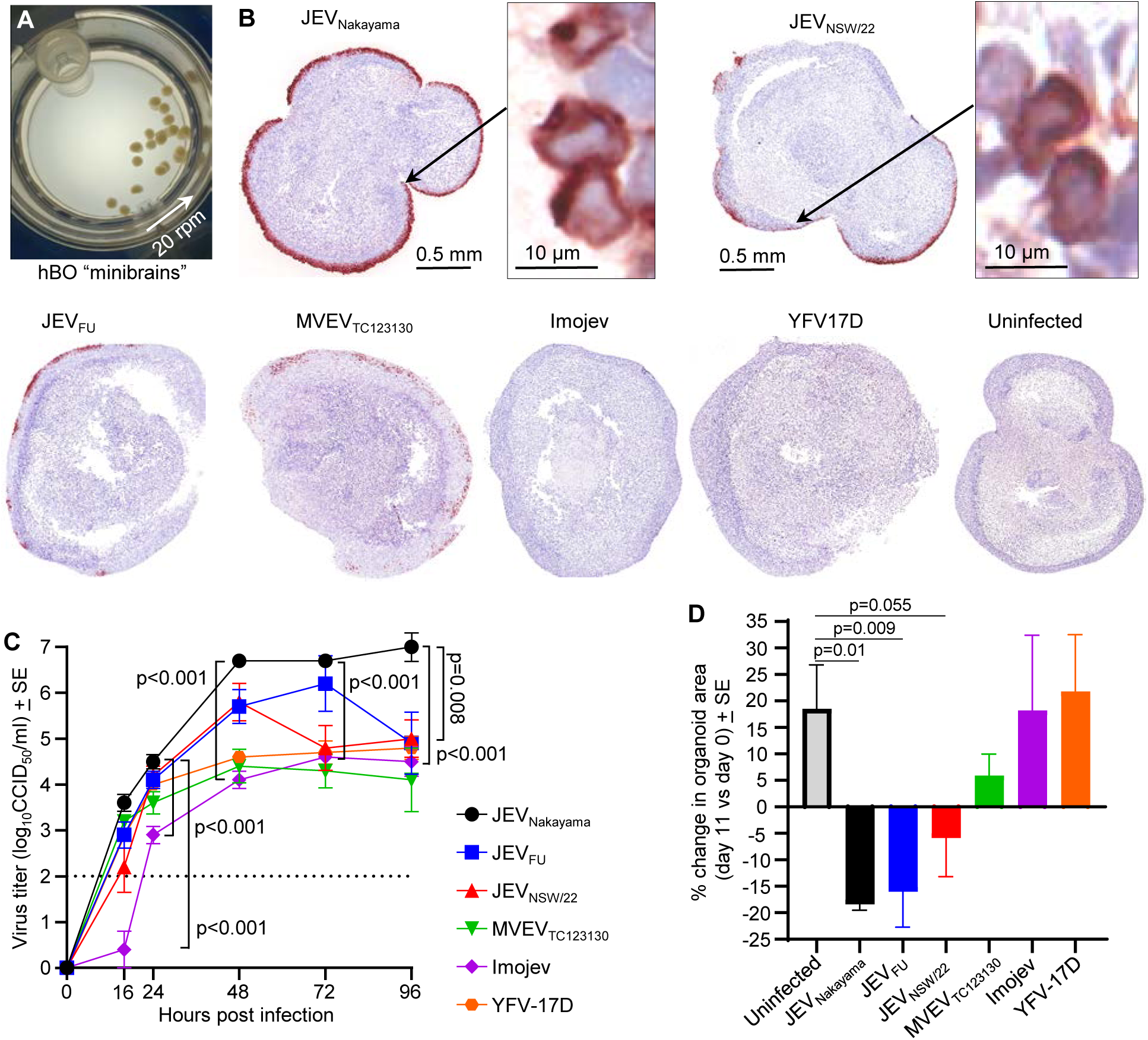
Infection of human cortical brain organoids (hBOs). (A) Photograph of “mini-brains” cultured in a rotating CelVivo Clinostar incubator. (B) IHC of viral antigen (4G4) for hBOs at 4 dpi. Images are representative of n=4 hBOs for each group. Magnified images of sparse Imojev and YFV 17D infected cells are shown in Supplementary Figure 12A. (B) Viral growth kinetics up to 4 dpi determined by CCID_50_ assays of culture supernatants at the indicated hours post infection; limit of detection is 2 log_10_CCID_50_/ml. At all time points JEV_Nakayama_ vs. Imojev, and at 96 h JEV_Nakayama_ vs. JEV_NSW/22_ were significant (t tests, n=5 organoids per group). (C) Mean percentage change in organoid area at 11 dpi vs. 9 dpi for each organoid (n=8 for uninfected and MVEV_TC123130_, otherwise n=4). Statistics are by Kolmogorov-Smirnov test for uninfected versus JEV_Nakayama_, and t-test for uninfected versus JEV_FU_ or JEV_NSW/22_.

All viruses were able to productively infect hBOs, with JEV_Nakayama_ infection generating higher viral titers than Imojev at all time points (Figure 7C, p<0.001). By 96 h JEV_Nakayama_ titers were also significantly higher than those seen for JEV_NSW/22_ (Figure 7C).

Over 11 days, uninfected hBOs grew in circumference by ∼20% (Figure 7D, Uninfected). hBOs infected with JEV isolates shrank in circumference by ∼5% to 15% by 11 dpi, although when compared with uninfected organoids, this only approached significance for JEV_NSW/22_ (Figure 7D; Supplementary Figure 13B-C). hBOs infected with MVEV_TC123130_, Imojev and YFV 17D infected hBO did not shrink significantly when compared with uninfected controls, although the MVEV_TC123130_ data suggested a marginal reduction in circumference (Figure 7C; Supplementary Figure 13C). These data (Figure 7D) reflect differences in viral replication (Figure 7C) and/or viral protein immunohistochemistry (Figure 7B) and likely reflect virus-induced CPE.

All viruses replicated productively in a human neural progenitor cell line, RENcell VM, with all viruses except Imojev causing fulminant CPE (Supplementary Figure 14A,B). Although Vero E6 cells are widely used for flavivirus research, CPE induced by JEV_NSW/22_ was considerably less pronounced in these cells (Supplementary Figure 14C).

### Human post Imojev-vaccination sera neutralizes JEV and MVEV with titers related to envelope protein amino acid conservation

Imojev is one of two JEV vaccines available in Australia. The Imojev prME genes are derived from the genotype 3 JEV_SA14-14-2_ (33, 64) strain, which was attenuated via extensive *in vitro* and *in vivo* passaging (Supplementary Figure 1). Most flavivirus neutralizing antibodies recognize epitopes on the envelope protein, particularly in the putative receptor binding domain III (67). JEV_Nakayama_ and the JEV_SA14-14-2_ component of the Imojev vaccine both belong to genotype 3, but JEV_Nakayama_ has 96.8% envelope protein amino acid identity to Imojev (Figure 8A). JEV_FU_ has 96.4% envelope protein identity, while JEV_NSW/22_ has drifted further from the genotype 2 and 3 strains with 93.4% envelope protein identity (Figure 8A). MVEV_TC123130_, which is the closest phylogenetically related flavivirus to JEV (Supplementary Figure 1B), has 80.4% envelope amino acid identity (Figure 1A). Alignment of the envelope amino acid differences for these strains compared to Imojev reveal that a disproportionate number of the non-conservative changes were in domain III (Figure 8A).

**Figure 8.**
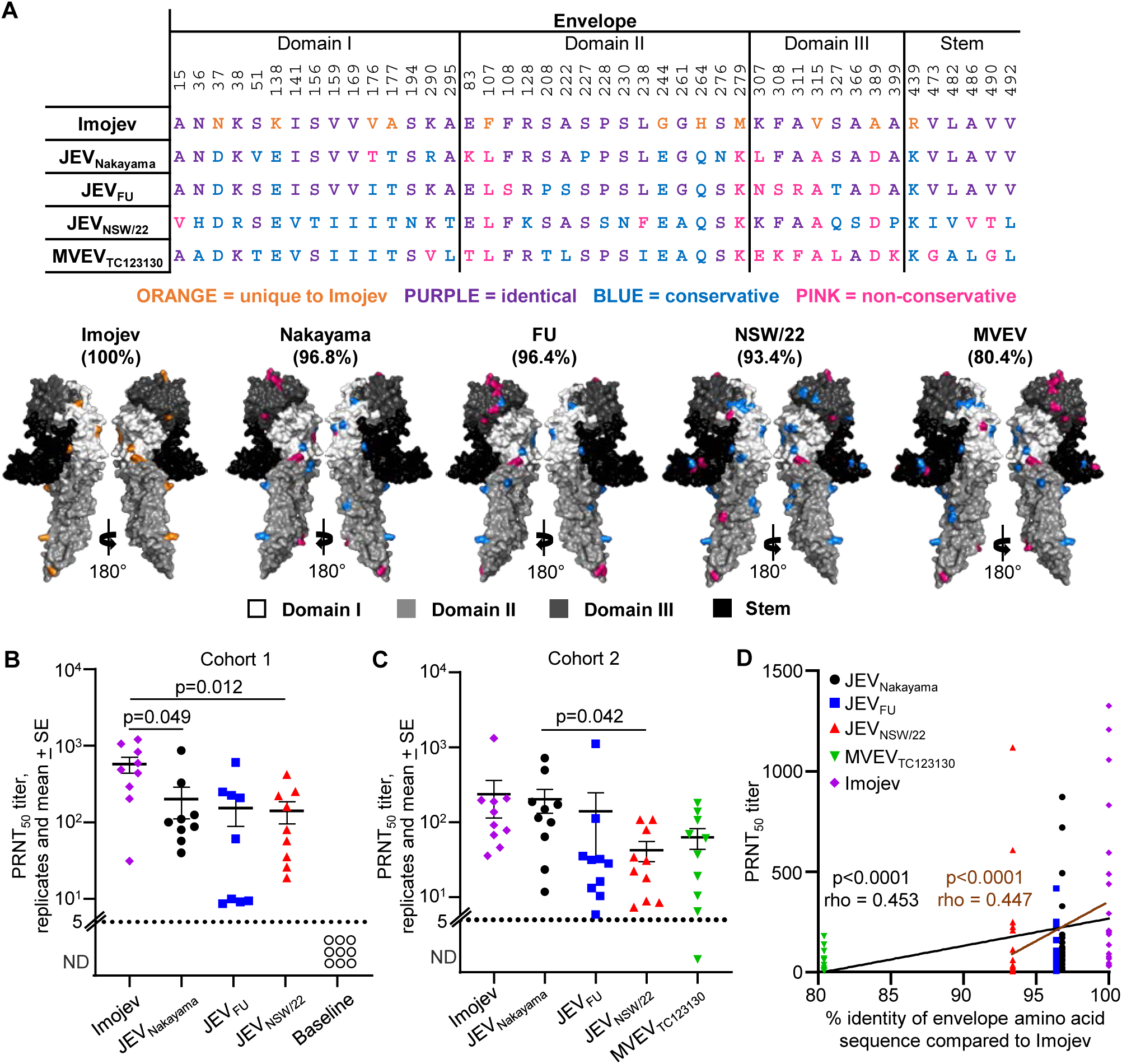
Human post Imojev-vaccination sera neutralizes JEV and MVEV with titers related to envelope protein amino acid conservation. (A) Envelope protein (domains I, II, III and STEM) amino acid sequences for Imojev, JEV_Nakayama_, JEV_FU_, JEV_NSW/22_, and MVEV_TC123130_ (refer to Supplementary Figure 1 for GenBank accession numbers). Sequences for isolates were aligned using MEGA-X and the ClustalW plugin with default parameters. Colouring indicates amino acid category compared to Imojev (orange = unique to Imojev, purple = identical, blue = conservative amino acid difference, pink = non-conservative amino acid difference (38)). Crystal structure of JEV envelope (PDB: 5WSN) with amino acid differences for JEV_Nakayama_, JEV_FU_, JEV_NSW/22_, and MVEV_TC123130_ compared to Imojev coloured as described in the table. Percentages indicate percent sequence identity relative to Imojev. (B) Human serum taken at day 0 and day 28 post-Imojev vaccination (n=9, cohort 1) was used in plaque reduction neutralisation assays against JEV_Nakayama_, JEV_FU_ and JEV_NSW/22_, and the plaque reduction neutralisation 50% titer (PRNT_50_) was calculated. Mean and standard error are shown. Statistics are paired t-test comparing Imojev with JEV_Nakayama_ or JEV_NSW/22_. (B) Human serum taken 2-12 months post-Imojev vaccination (n=10, cohort 2) was used in plaque reduction neutralisation assays against JEV_Nakayama_, JEV_FU_, JEV_NSW/22_, and MVEV_TC123130_ and the plaque reduction neutralisation 50% titer (PRNT_50_) was calculated. Mean and standard error are shown. Statistics are paired t-test comparing JEV_Nakayama_ with JEV_NSW/22_. (D) PRNT_50_ titers in ‘B’ and ‘C’ plotted against percentage envelope protein amino acid identity in ‘A’. Curve fit is shown, and statistics calculated by Spearman correlation with p and rho values shown (black line represents all data, brown line excludes MVEV_TC123130_ from analysis).

Serum neutralizing antibodies post-vaccination is currently viewed as the best measurable correlate of vaccine protection for JEV (68). To determine if the Australian outbreak genotype 4 JEV_NSW/22_ is neutralized by antibodies produced in response to Imojev vaccination, serum was taken from two human cohorts. Cohort 1 was serum collected from individuals at approximately 28 days post-vaccination, while cohort 2 serum was collected at variable times post-vaccination (2 months to 1 year). Ethical approvals for cohort 1 serum allowed for neutralisation assays against the Imojev vaccine itself, JEV_Nakayama_, JEV_FU_, and JEV_NSW/22_ (Figure 8B), while ethical approval for cohort 2 additionally allowed for neutralisation assays against MVEV_TC123130_ (Figure 8C). All serum samples from both cohorts had a measurable 50% neutralisation titer (PRNT_50_) titer against JEV_Nakayama_, JEV_FU_, and JEV_NSW/22_ (Figure 8B-C), indicating that seroconversion provided neutralizing antibodies that cross-react between JEV genotypes 2, 3 and 4. In cohort 1, PRNT_50_ titers were significantly lower against JEV_Nakayama_ and JEV_NSW/22_ compared to against Imojev (Figure 8A), suggesting key antigenic differences from the vaccine. In cohort 2, PRNT_50_ titers against JEV_Nakayama_ were not significantly different compared to Imojev, however PRNT_50_ titers were significantly lower for JEV_NSW/22_ (Figure 8C). Similar conclusions were drawn when the raw percentage of plaque neutralisation at a high serum dilution (1:160) was compared between virus strains (Supplementary Figure 15). With PRNT_50_ data from both cohorts combined, the percentage amino acid identity of the envelope protein compared to the Imojev vaccine significantly correlated with PRNT_50_ titers (Figure 8D, black line). The significance (p-value) and the correlation coefficient (rho), were similar when the same analysis was conducted excluding the PRNT_50_ data for MVEV_TC123130_ (Figure 8D, brown line). Overall, our data indicates that Imojev vaccination provided neutralizing antibodies against JEV_NSW/22_ in all individuals, but the level of cross-neutralisation were related to the conservation in envelope protein amino acid sequences.

## Discussion

Herein we provide a comprehensive *in vivo* and *in vitro* characterization of the genotype 4 JEV_NSW/22_ isolate from the recent Australian outbreak, and illustrate mouse models of infection and rare CNS neuropathological manifestations that recapitulate many aspects of human and primates disease (6, 44–47, 69). The capacity of JEV_NSW/22_ to cause lethal neuroinvasive infection in mice was significantly diminished compared to JEV_Nakayama_ and JEV_FU_, with only one *Irf7^-/-^*mouse succumbing to JEV_NSW/22_ infection out of 63 infected C57BL/6J or *Irf7^-/-^* mice. Such rare lethal neuroinvasion recapitulates what is seen in humans, with approximately 1 in 750 infections causing fatality (2, 3). Serosurvey data, albeit limited, suggests the ratio of human symptomatic to asymptomatic infections is not particularly different for JEV_NSW/22_. To date 45 clinical cases have been notified for the recent outbreak in Australia (70), with serosurveys in Victoria (n=820 participants) and New South Wales (NSW) (n=917 participants) reporting 3.3% and 8.7% of participants as seropositive for JEV, respectively (71, 72). Although somewhat dated, population data for the primary recruitment locations for the serosurveys is available from Australian Bureau of Statistics 2016, with Victorian recruitment locations providing a population total of 160,294 (Mildura, Lockington, Shepparton, Cobram, Yarrawonga, Rutherglen, Wodonga, Wangaratta, Rochester), and NSW locations a total of 68,431 (Balranald, Corowa, Dubbo, Griffith, Temora). As [0.033 x 160,294] + [0.087 x 68,431] / 250 = 45, the serosurvey data is consistent with the expected symptomatic to asymptomatic ratio of ≈1 in 250 for JEV and thus provides no compelling evidence for overt virulence differences for JEV_NSW/22_ in human populations.

Our *Irf7^-/-^* mouse model of JEV_NSW/22_ provides for a more robust viremia, and a slightly higher chance of lethal neuroinvasive infection. Increased lethal neuro-penetrance in *Irf7^-/-^* mice was associated with a prolonged viremia, possibly via increased inflammation-driven blood brain barrier breakdown as a result (5, 73). The use of *Irf7^-/-^* mice to increase lethal neuroinvasive infection compared to C57BL/6J was only suitable for JEV_FU_ and JEV_NSW/22_. This was likely due to higher sensitivity to type I IFN for these isolates, demonstrated using MEF cells, compared to JEV_Nakayama_ and MVEV_TC123130_. The partially defective type I IFN responses in *Irf7^-/-^* mice (24) thus provides a benefit for JEV_FU_ and JEV_NSW/22_ neuro-penetrance, but not for JEV_Nakayama_ or MVEV_TC123130_. When type I IFN responses were completely defective (*Ifnar^-/-^* mice and *Irf3/7^-/-^*MEFs), differences between virus replication and/or lethality were minimal. Overall, these results suggest that JEV_NSW/22_ may be more sensitive to, or less able to suppress, type I IFN responses. Inhibition of type I IFN responses is mediated by NS5 for inhibition of STAT2 and NS3 (74) and subgenomic flavivirus RNA (sfRNA)/NS5 (21) for inhibition of STAT1. Although the latter likely operates for WNV in mice (75) and is involved in promoting apoptosis (21), the efficiency of these systems during JEV infection of humans and mice remains to be determined. sfRNA is derived from the 3’UTR (21), where JEV_NSW/22_ does show a small number of nucleic acid changes, but which are unlikely to affect sfRNA production (Supplementary Table 3). Mouse models of flavivirus pathogenesis frequently use *Ifnar^-/-^* mice (30, 76, 77) as flaviviruses often replicate poorly in wild-type mice as the ability of NS5 to suppress the antiviral type I IFN responses in humans often fails to operate in mice (78). We show herein that *Ifnar^-/-^*mice are not a good model for JEV neuropathology, as they reach ethically defined endpoints before brain infection can occur. However *Ifnar^-/-^* mice are a good model of robust and lethal viremia, which may provide a useful and stringent model for vaccine testing.

JEV_NSW/22_ does not contain known attenuating mutations that would explain its reduced virulence in mice. There are 9 amino acids in the Imojev vaccine envelope gene that have been associated with attenuated neurovirulence; F107, K138, V176, A177, G244, H264, M279, V315 and R439 (79–89). At these positions, JEV_Nakayama_, JEV_FU_, JEV_NSW/22_, and MVEV_TC123130_ all have the same amino acids (L107, E138, T177, E244, Q264, K279, A315, and K439), except for position 176 where JEV_Nakayama_ has T176, and JEV_FU_, JEV_NSW/22_, and MVEV_TC123130_ have I176 (Supplementary Table 3). JEV_NSW/22_ retains E at position 138, and this amino acid has been identified by several studies as a principal neurovirulence determinant (90–94), with a role in neuronal cell binding hypothesized (37). YFV 17D has a valine (V) at this residue, possibly contributing to reduced neurovirulence (95). NS1 and NS2A have been implicated in neuroinvasion, but not neurovirulence (88). Among the six changes in NS1 associated with attenuation of neuroinvasiveness are R147H and R339M (88), of which H147 is present in both JEV_Nakayama_ and JEV_NSW/22_, with K339 (a conserved substitution for R) present in JEV_NSW/22_.

JEV non-structural protein 4B (NS4B) alone can induce apoptosis and encephalitis (96), however, NS4B is completely conserved between JEV_Nakayama_, JEV_FU_, and JEV_NSW/22_ (Supplementary Table 3). prM has been reported to influence neuroinvasiveness of ZIKV (76) and JEV virulence in mice (97). JEV_NSW/22_ has a number of unique changes in prM (Supplementary Table 3), although their functional implications remain unclear. JEV_NSW/22_ also has an additional N-linked glycosylation site at position 175 in NS1 that is lacking in JEV_Nakayama_ and JEV_FU_ (Supplementary Table 3). However, this N-linked glycosylation site is reported to increase WNV neuroinvasiveness in wild-type mice (98, 99), a trend not seen in our JEV data (Figure 1F). Thus JEV_NSW/22_ shows no obvious sequence characteristics that can be ready associated with the reduced virulence in mice, and mutagenesis experiments would be required to fully understand these differences.

All human participants vaccinated with the Imojev vaccine induced neutralizing antibodies to JEV_NSW/22_, suggesting that this vaccine, which is available in Australia, is likely to afford some protection against Australian outbreak genotype 4 JEV. However, divergence of envelope protein amino acid sequences from that of the Imojev vaccine affected the PRNT_50_ titers, although it is unclear how this may translate to the impact on vaccine efficacy, especially given that *in vitro* neutralisation assays do not capture the full range of protective mechanisms mobilised *in vivo* (100). Nonetheless, this provides a strong rationale for development of updated JEV vaccines that use antigen sequences from currently circulating JEV strains, such as genotype 4 in Australia (14), genotype 5 in Republic of Korea (12), and genotype 1 in most other areas of South East Asia (101). Imojev vaccination also produced neutralizing antibodies against MVEV_TC123130_ which is consistent with previous studies using other JEV vaccines (102, 103), and is consistent with cross-reactivity in serology-based diagnostic assays (13, 104). There is also some evidence that JEV vaccination or infection provides partial cross-protection against MVEV and vice versa (105–107).

Although one limitation of this study may be that JEV_NSW/22_ was isolated from a pig, there are only 4 amino acid differences between this isolate and a JEV G4 sequence from a human brain in the Tiwi Islands (Northern Territory, Australia) in 2021 (Genbank accession OM867669 (14)).

The differences are; envelope-238 F vs. L, NS2A-71 I vs. T, NS2B-59 E vs. G, NS3-436 E vs. G. In addition, JEV_Nakayama_ was passaged in suckling mouse brains, which may contribute to the increased neuroinvasion in C57BL/6J mice, although the adaptive mutations acquired during passaging, if any, are currently unknown. Furthermore, JEV_FU_ has not been passaged in mice, but was still more lethal than JEV_Nakayama_ in *Irf7^-/-^* mice. Use of C57BL/6J mice that lack a functional nicotinamide nucleotide transhydrogenase *(Nnt)* may be another issue, as background and *Nnt* are able to affect viral immunopathogenesis (25, 59). However, we found that neither *Nnt* nor a C57BL/6N genetic background significantly impacted JEV replication or immunopathology (Supplementary Figure 16).

In conclusion, we show that JEV_NSW/22_ has reduced neuropenetrance in mice but retains capacity for rare lethal neuroinvasion, consistent with reported human fatalities in the 2022 Australian outbreak.

## AUTHOR CONTRIBUTIONS

Conceptualization, D.J.R. Methodology, D.J.R., W.N., A.S, G.D., N.G., and N.W. Formal analysis, W.N., D.J.R. and A.S. Investigation, W.N., B.T., R.S., A.C., K.Y., N.G., T.L., and D.J.R. Resources, R.S., N.G., G.D., N.W., and A.S. Data curation, W.N., D.J.R., N.G., and A.S. Writing, original draft – D.J.R. Writing, review and editing, A.S and W.N. Visualization, D.J.R., W.N. and A.S. Supervision, D.J.R., G.D. and A.S. Project administration, D.J.R., G.D., N.W., and A.S. Funding acquisition, A.S., D.J.R, and G.D, and N.W.

## ACKNOWLEDGEMENTS

From QIMR Berghofer MRI we thank Dr I Anraku for managing the PC3 (BSL3) facility, animal house staff for mouse breeding and agistment, Drs Anthony White and Lotta Oikari for providing the iPSC line, Crystal Chang, Sang-Hee Park and Ashwini Potadar for the histology and immunohistochemistry, Dr Viviana Lutzky for proof reading, and Dr. Gunter Hartel (Head of Statistics, QIMRB) for assistance with statistics. We thank Dr. Peter Kirkland for supplying the JEV_NSW/22_ isolate. We thank Dr. Andrii Slonchak for insights into whether 3’UTR changes may affect sfRNA production.

## DATA AVAILABILITY

All data is provided in the manuscript and accompanying supplementary files.

## FIGURE LEGENDS

**Supplementary Figure 1.**
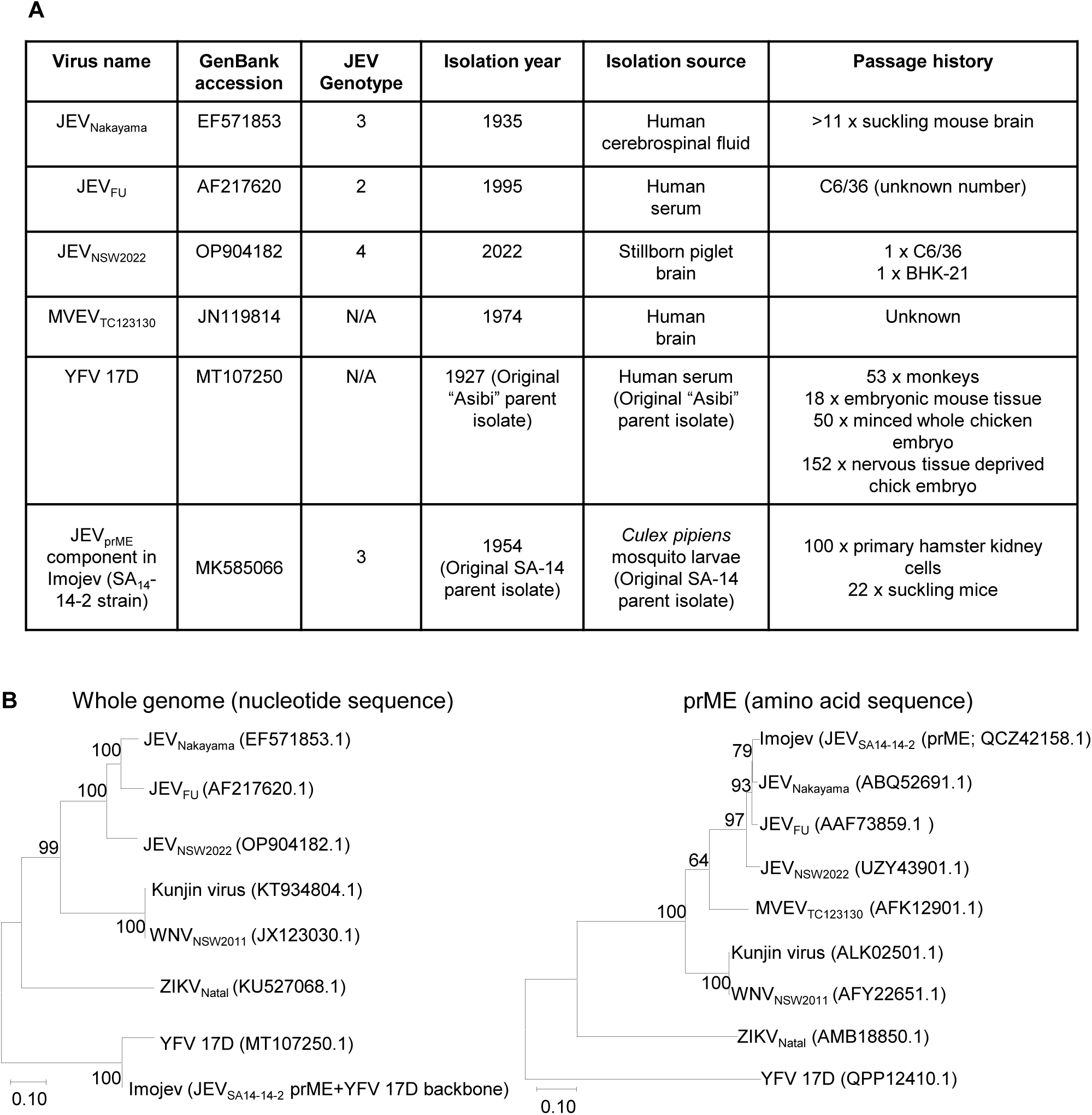
(A) Summary of virus isolates used in the study. (B) Phylogenetic tree for whole genome nucleotide sequence (left) and prME amino acid sequence (right). Phylogenetic trees were constructed after nucleotide or amino acid sequence alignment using MEGA-X (Molecular Evolutionary Genetics Analysis 10, Penn State University, State College, PA, USA) and the ClustalW plugin with default parameters. The phylogenetic tree was constructed using the Maximum Likelihood method and the General Time Reversible model (nucleotide sequence) or JTT matrix-based model (amino acid sequence). Whole genome sequence is not available for MVEV_TC123130_. Whole genome sequence for Imojev was constructed by combining the prME sequence from JEV_SA14-14-2_ with the remainder of the genome from YFV 17D.

**Supplementary Figure 2.**
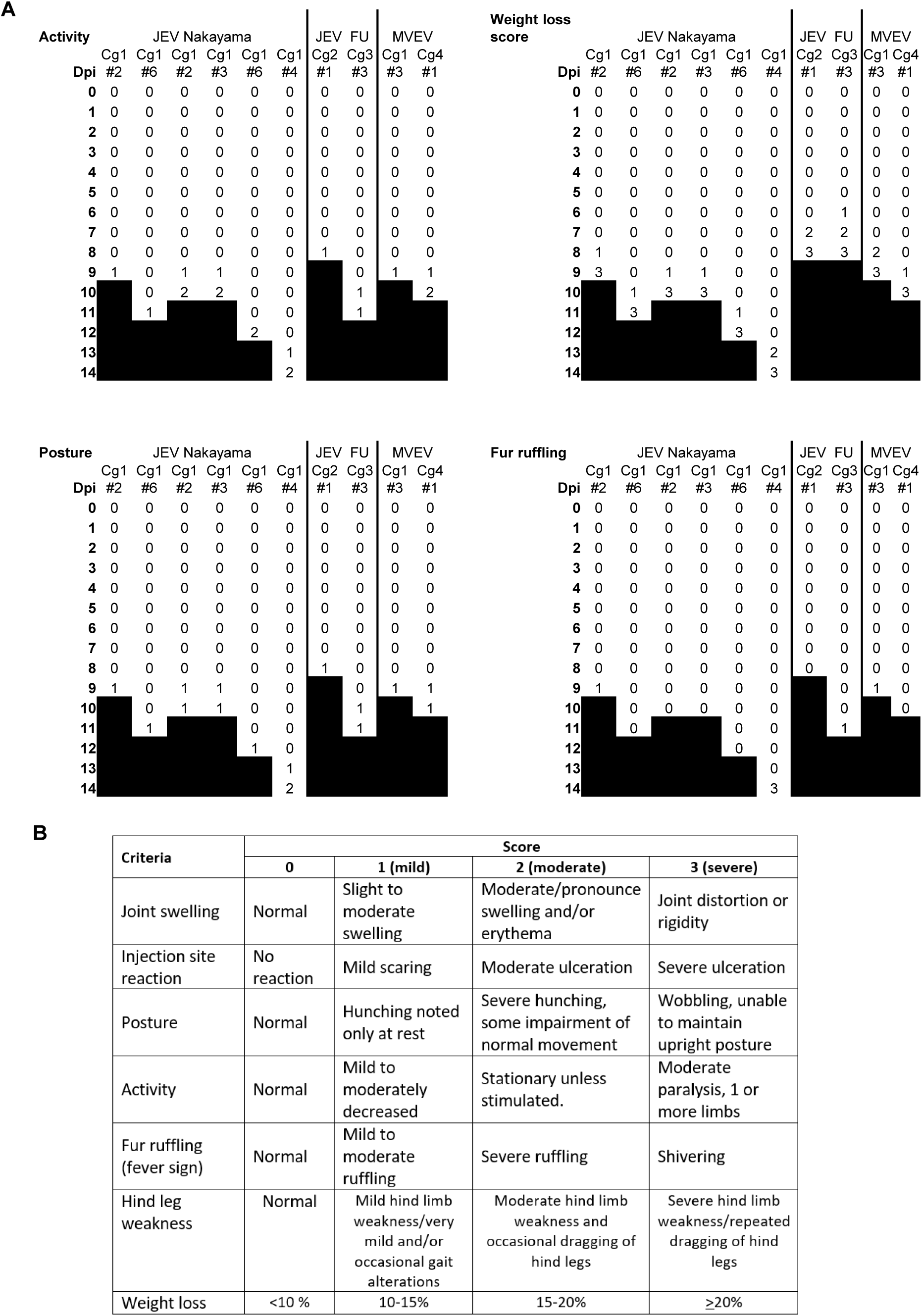
Disease scores for the ten C57BL/6J mice that were euthanized. (A) For the mice shown in Fig. 1F and I that were euthanized, the disease scores are shown. (B) The mice were monitored daily using the score card. Any animal reaching a level of 3 in any single criteria were euthanized. If an animal reaches a grade of 2 in two or more criteria the animal will be euthanized.

**Supplementary Figure 3.**
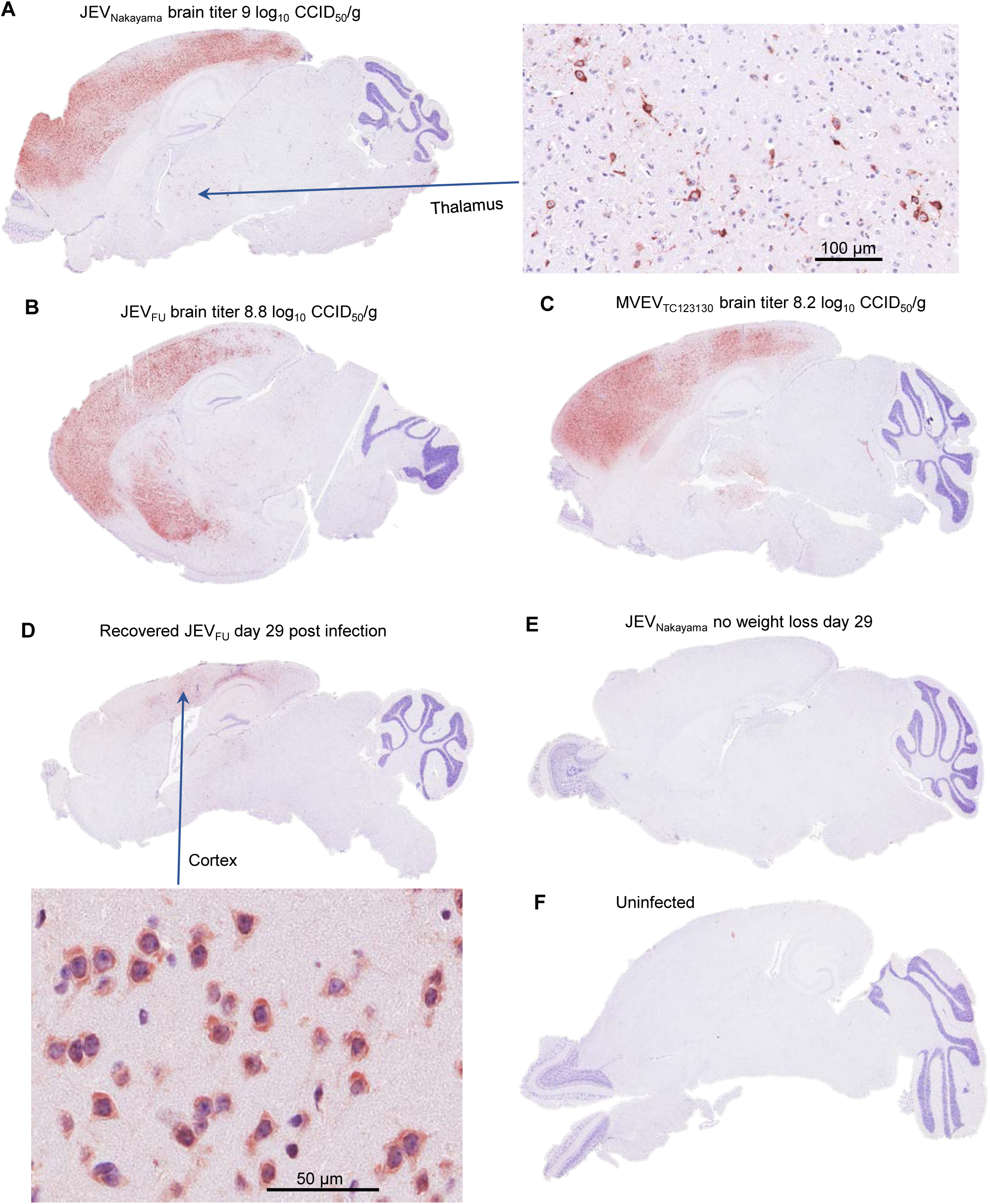
Viral antigen (NS1) staining in C57BL/6J mouse brains. IHC using a pan-flavivirus NS1 monoclonal antibody (4G4). (A-C) The brains from the 3 other C57BL/6J mice that required euthanasia in Fig. 1 (the fourth is shown in Fig. 2). (D) JEV_FU_ infected C57BL/6J mice that lost ∼15% body weight then recovered (from Fig. 1E). (E) A representative image of a mouse brain where infection did not lead to significant weight loss. (F) Uninfected control.

**Supplementary Figure 4.**
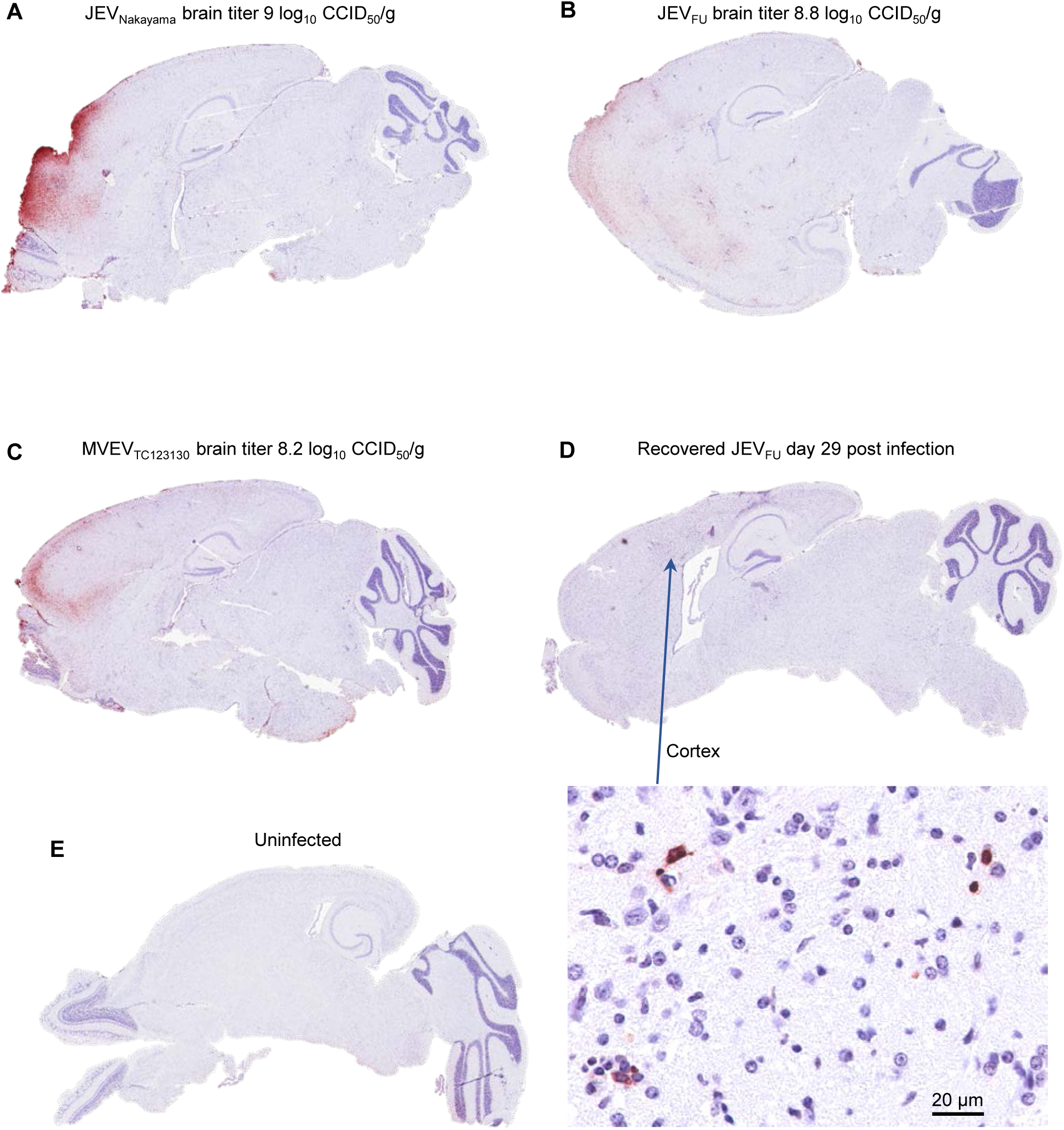
ApopTag staining in C57Bl/6J mouse brains. Staining for apoptosis using Apoptag. (A-C) The brains from the 3 other C57BL/6J mice that required euthanasia in Figure 1. (D) The JEV_FU_ infected C57BL/6J mice that lost ∼15% body weight then recovered (see Figure 1E). (E) Uninfected control.

**Supplementary Figure 5.**
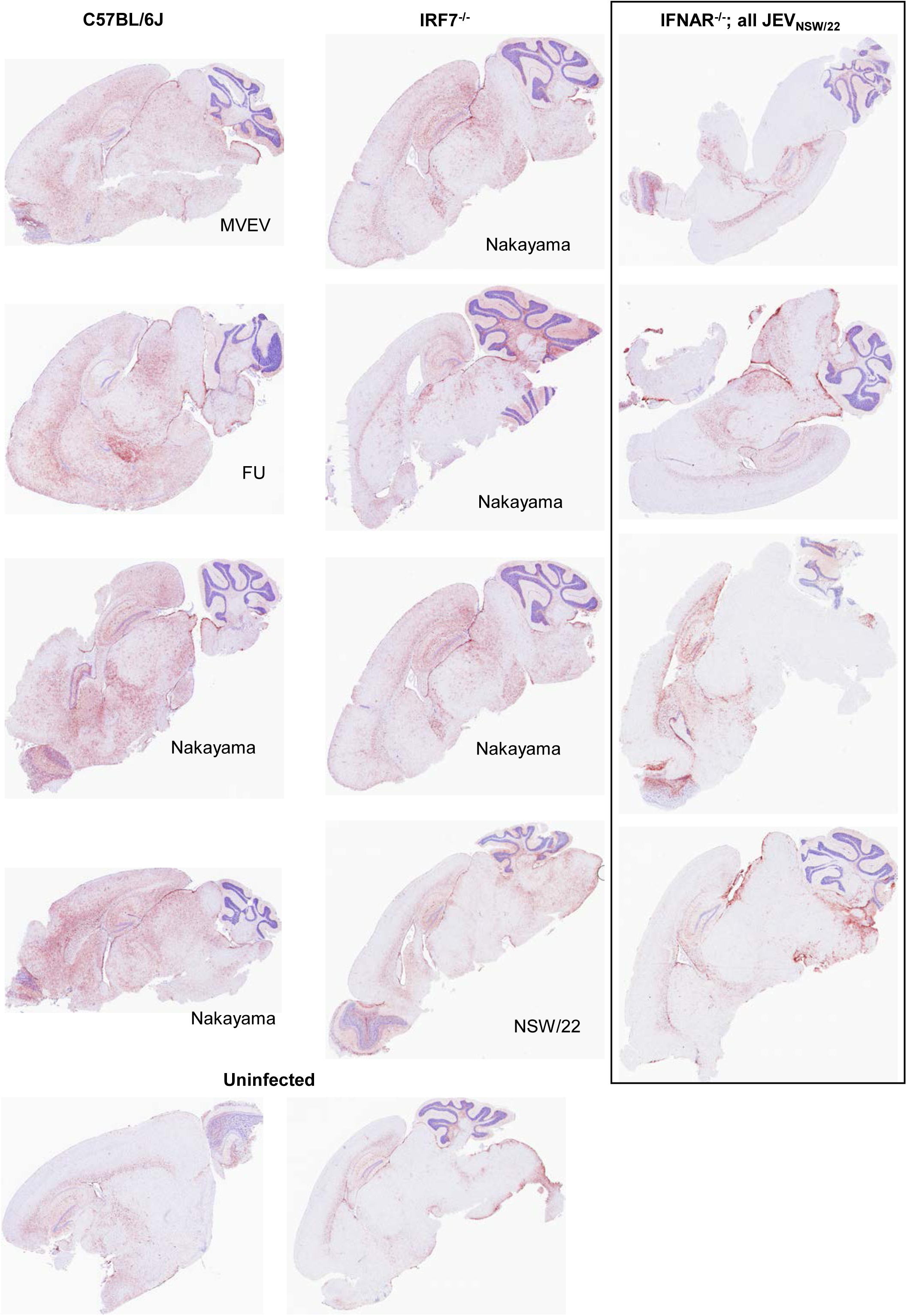
GFAP IHC for reactive astrocytes in mouse brains. Staining for reactive actrocyte using GFAP for the brains from C57BL/6J, IRF7^-/-^ and IFNAR^-/-^ mice that succumbed to the indicated virus. Uninfected mouse brains are also shown.

**Supplementary Figure 6.**
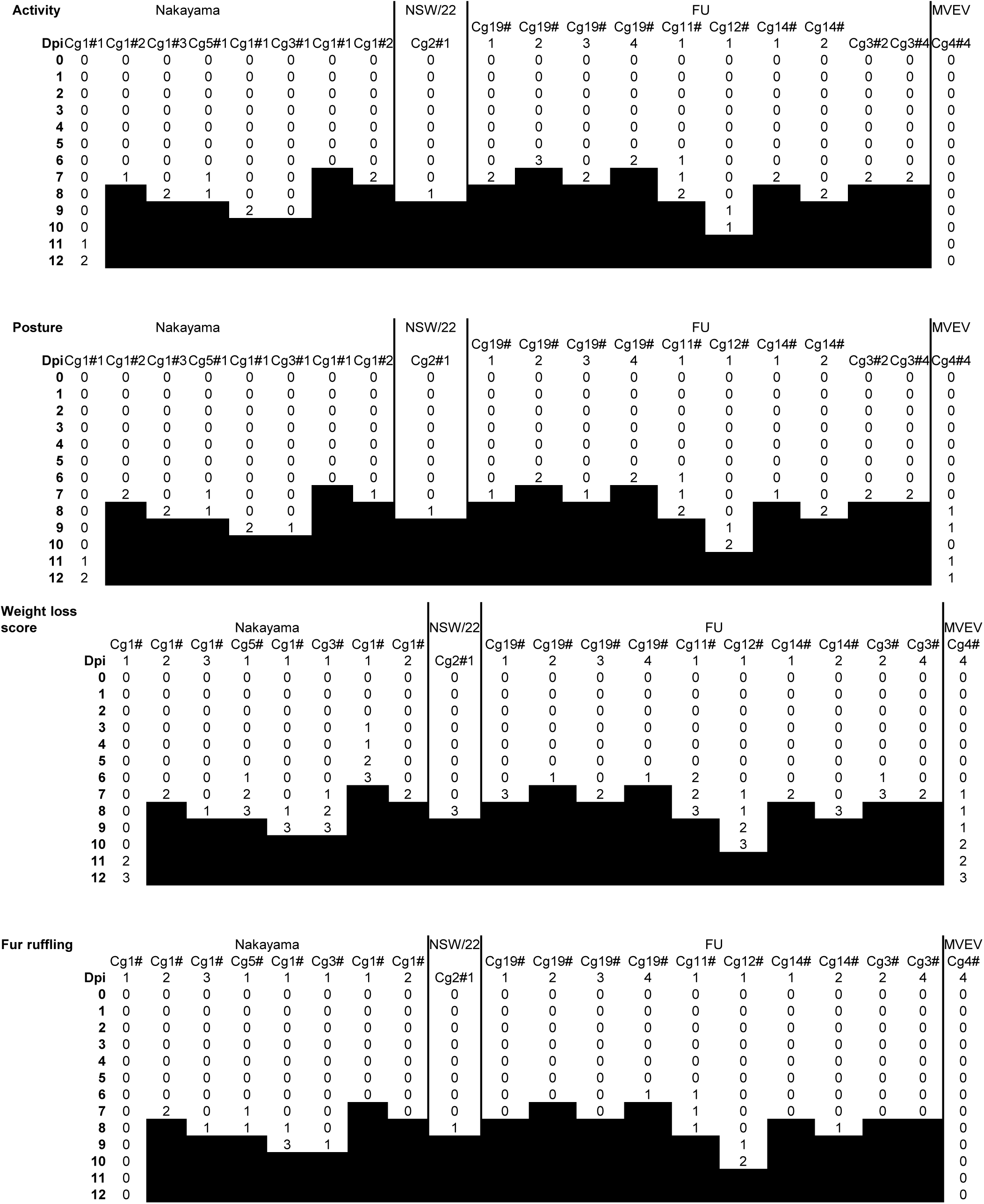
Disease scores for IRF7-/– mice that were euthanized. Mice described in Figure 3C, D. Scoring system shown in Supplementary Figure 2B.

**Supplementary Figure 7.**
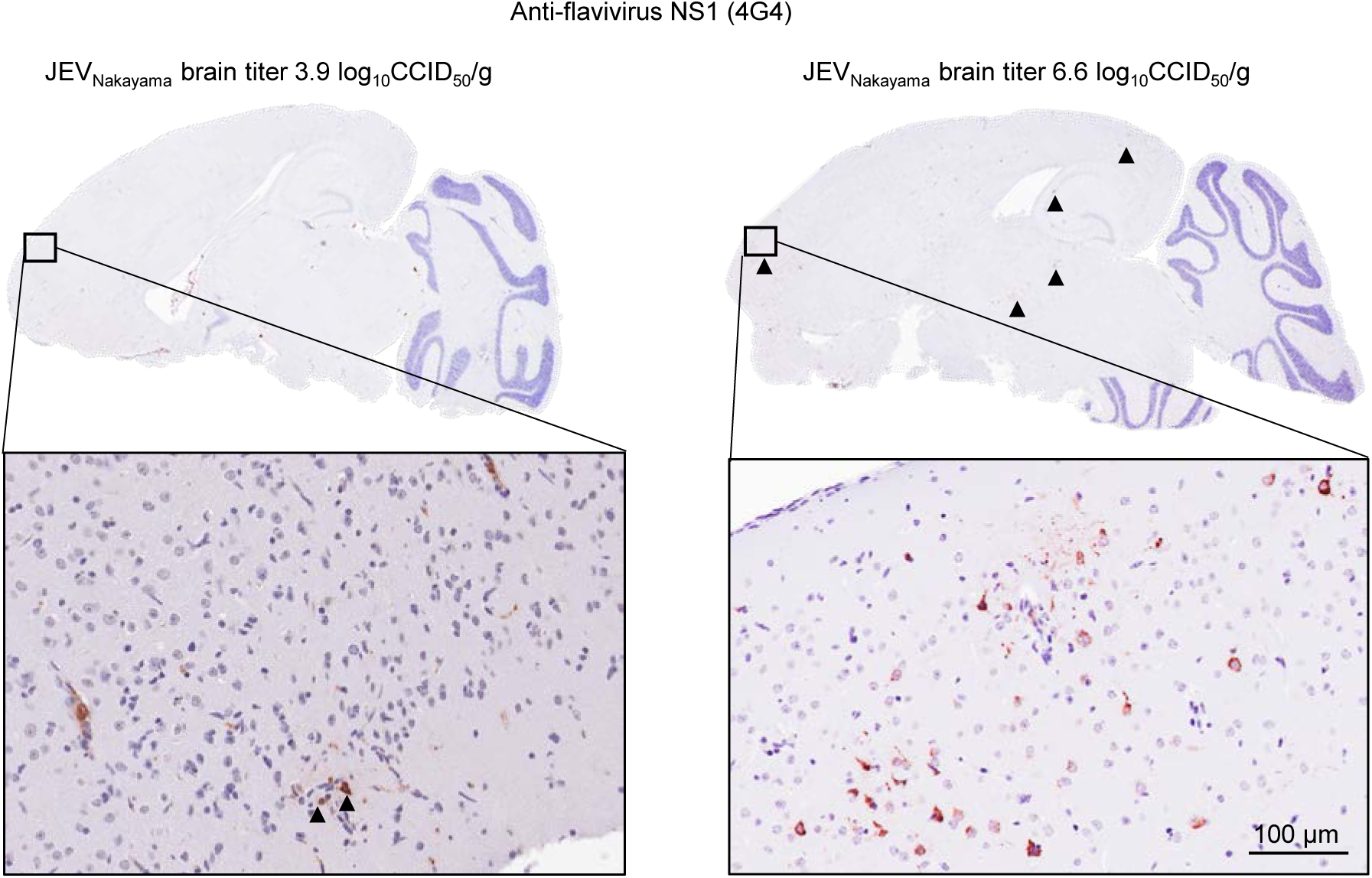
Viral antigen staining in IRF7^-/-^ mouse brains. IHC using a pan-flavivirus NS1 monoclonal antibody (4G4) for the brains from 2 other IRF7^-/-^ mice that required euthanasia (Figure 3). Low levels of staining of neurons were found in the JEV_Nakayama_ infected mice that had a low virus titer in the brain (3.9 log_10_CCID_50_/g) (left). Black arrowheads show small patches of staining in the cortex, hippocampus and thalamus (right).

**Supplementary Figure 8.**
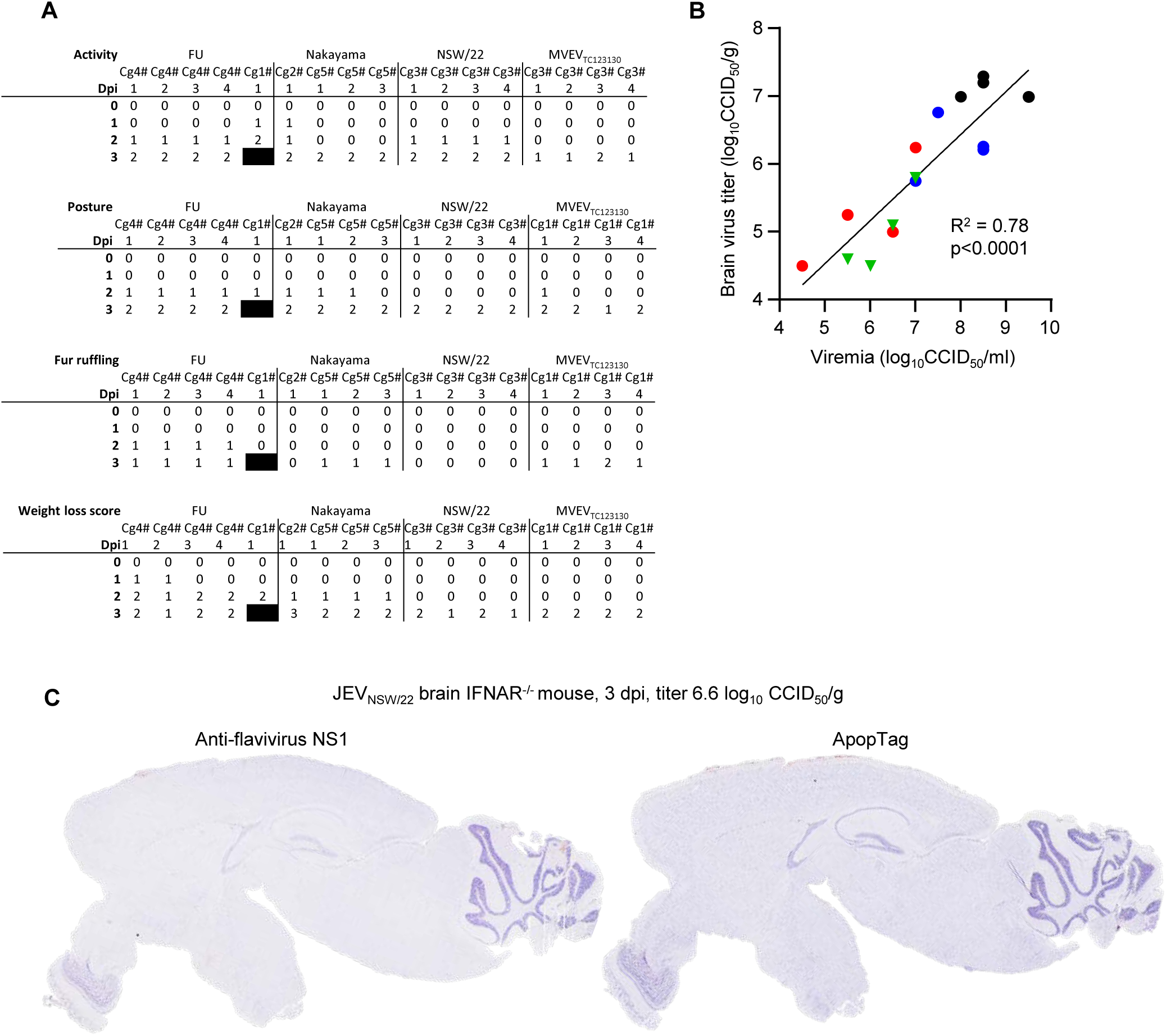
JEV and MVEV infection of IFNAR^-/-^ mice. (A) Disease scores as per Supplementary Figure 2B for mice shown in Figure 3. (B) Pearson correlation between brain titer (y-axis) from Figure 3E and viremia (x-axis) from Figure 3A. (B) IHC for flavivirus NS1 (left) or ApopTag (right) showing no detectable staining of brain cells. Images are representative of all IFNAR^-/-^ brains and JEV isolates.

**Supplementary Figure 9.**
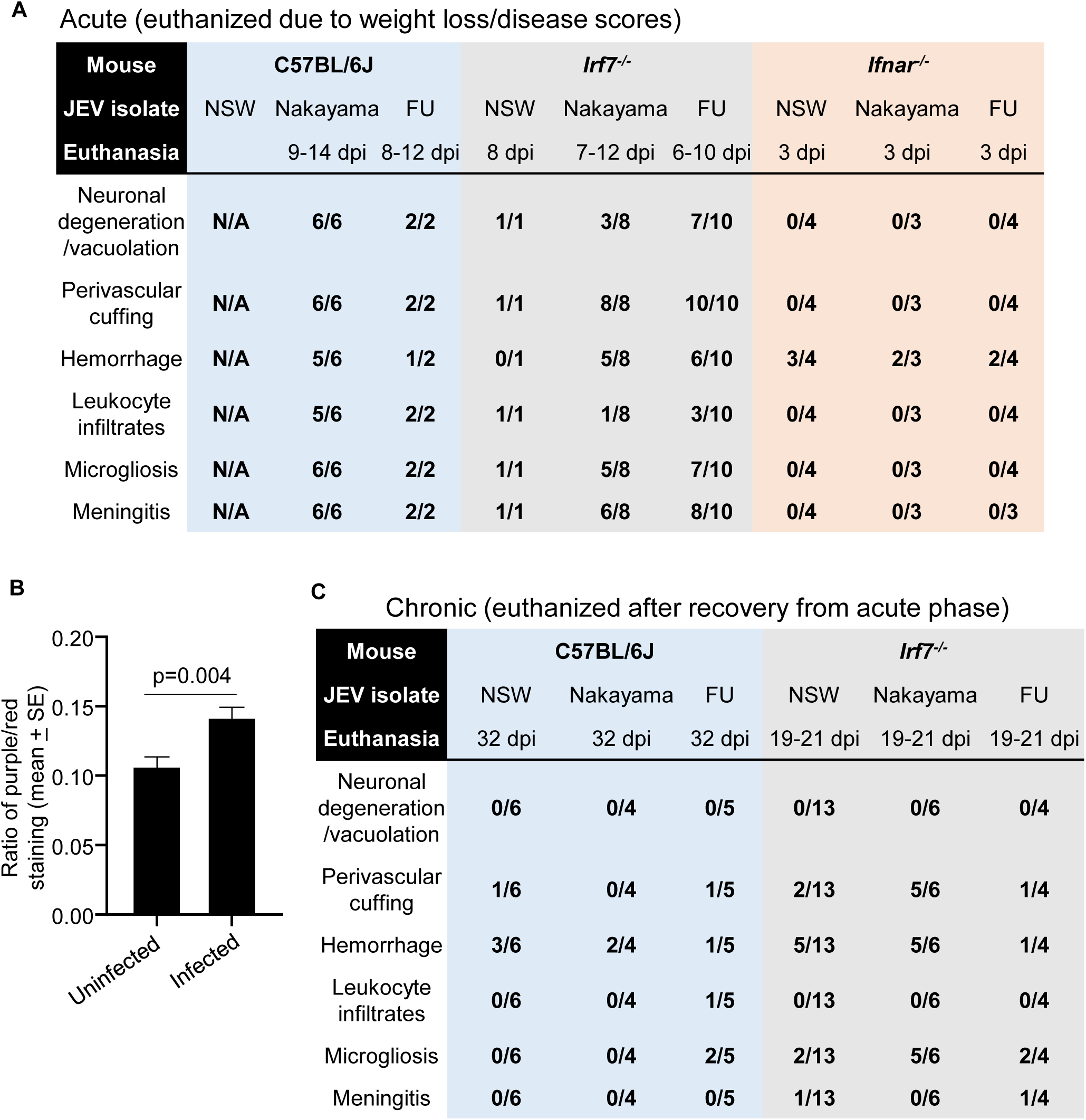
Lesion presence in H&E brain sections. A) Scoring for lesions described in Figure 6 for mice with acute disease. H&E scoring of 0 indicates no overt presence of these lesions. N/A – not available. B) Ratio of nuclear (blue/dark purple) to non-nuclear (red) staining of H&E stained brain sections (a measure of leukocyte infiltration). Data is the mean and standard error for n=30 infected and n=14 uninfected mouse brains from C57BL/6J and *Irf7^-/-^* mice that succumbed to JEV or MVEV infection (Fig. 1F, 1I and 3D). Statistics by t-test. C) As for ‘A’ but for mice that survived infection.

**Supplementary Figure 10.**
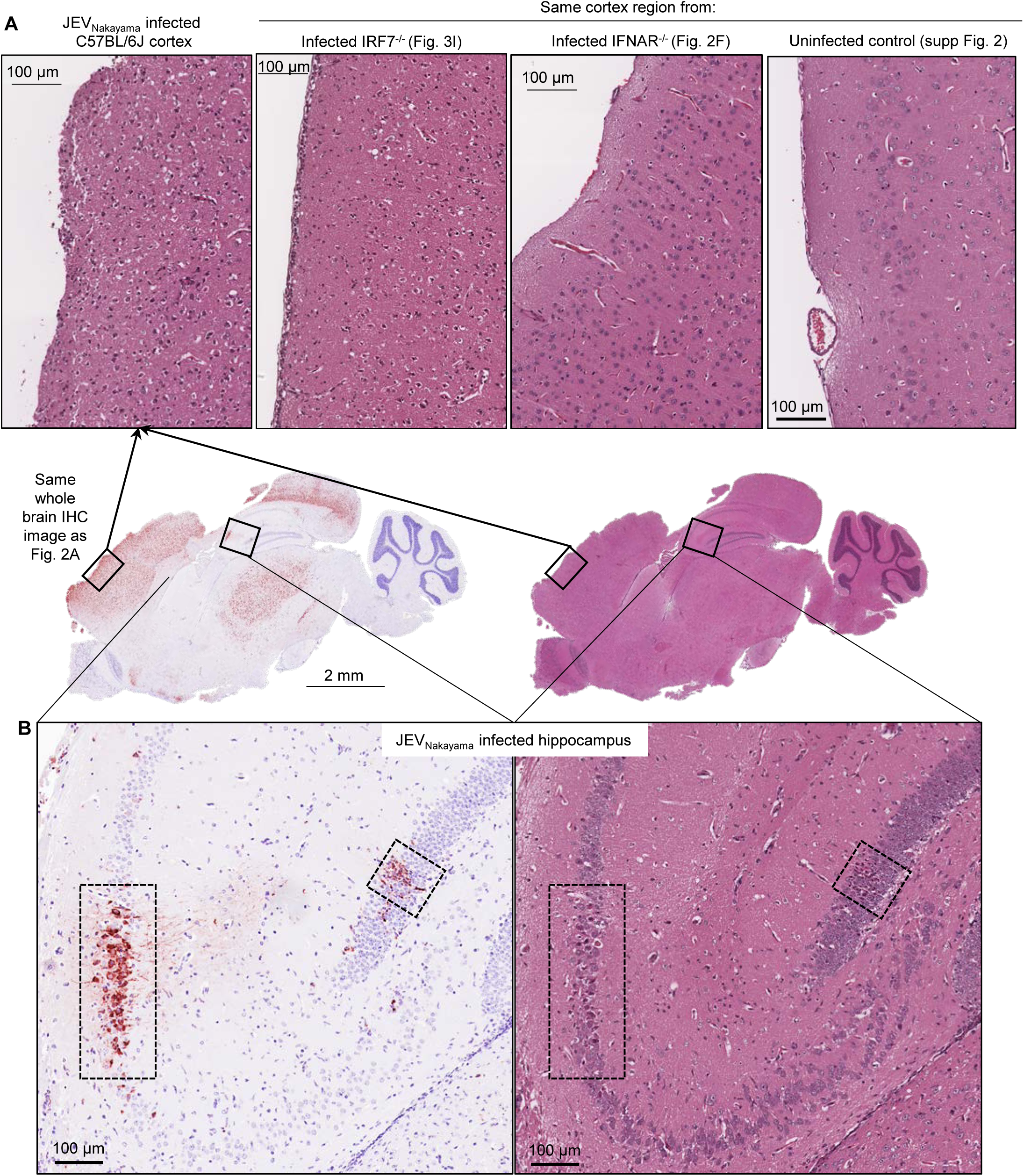
Histopathological lesions overlapped with areas of virus infection. (A) Cortex regions from JEV infected C57BL/6J, IRF7^-/-^, IFNAR^-/-^ mice or uninfected mice. The cortex regions from JEV infected mice were heavily infected, with most neurons stained positive for viral antigen. H&E detectable signs of neuron degeneration/vacuolation were concentrated in regions with high staining for viral antigen. (B) Viral antigen staining (left) in hippocampus overlapped with histological signs of neuron degeneration/vacuolation (right) (see Figure 6).

**Supplementary Figure 11.**
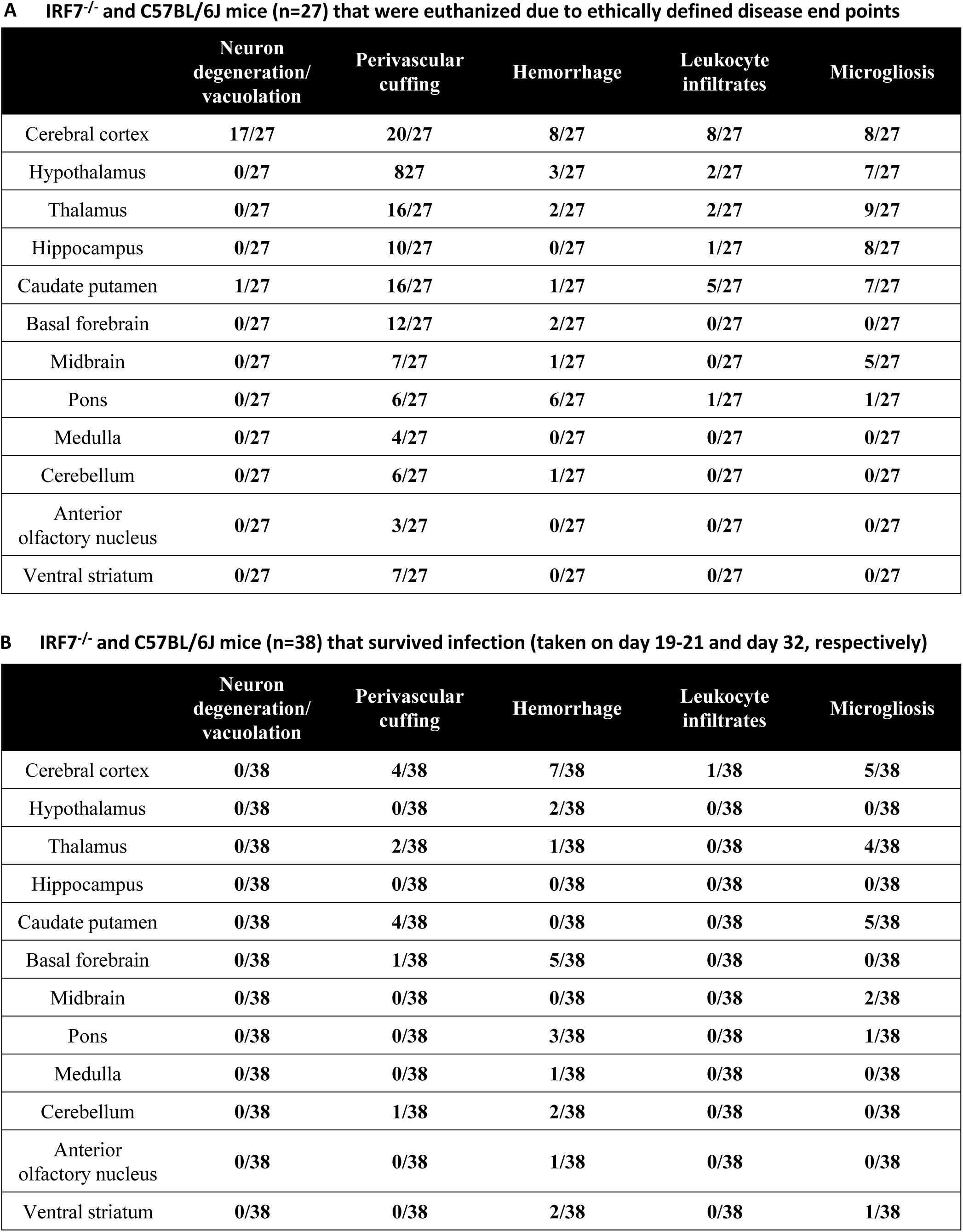
Histopathological lesions in specific brain regions for C57BL/6J and IRF7^-/-^. Scoring only reflects presence or absence of lesions, and does not indicate severity of lesions. (A) Acute (B) Chronic.

**Supplementary Figure 12.**
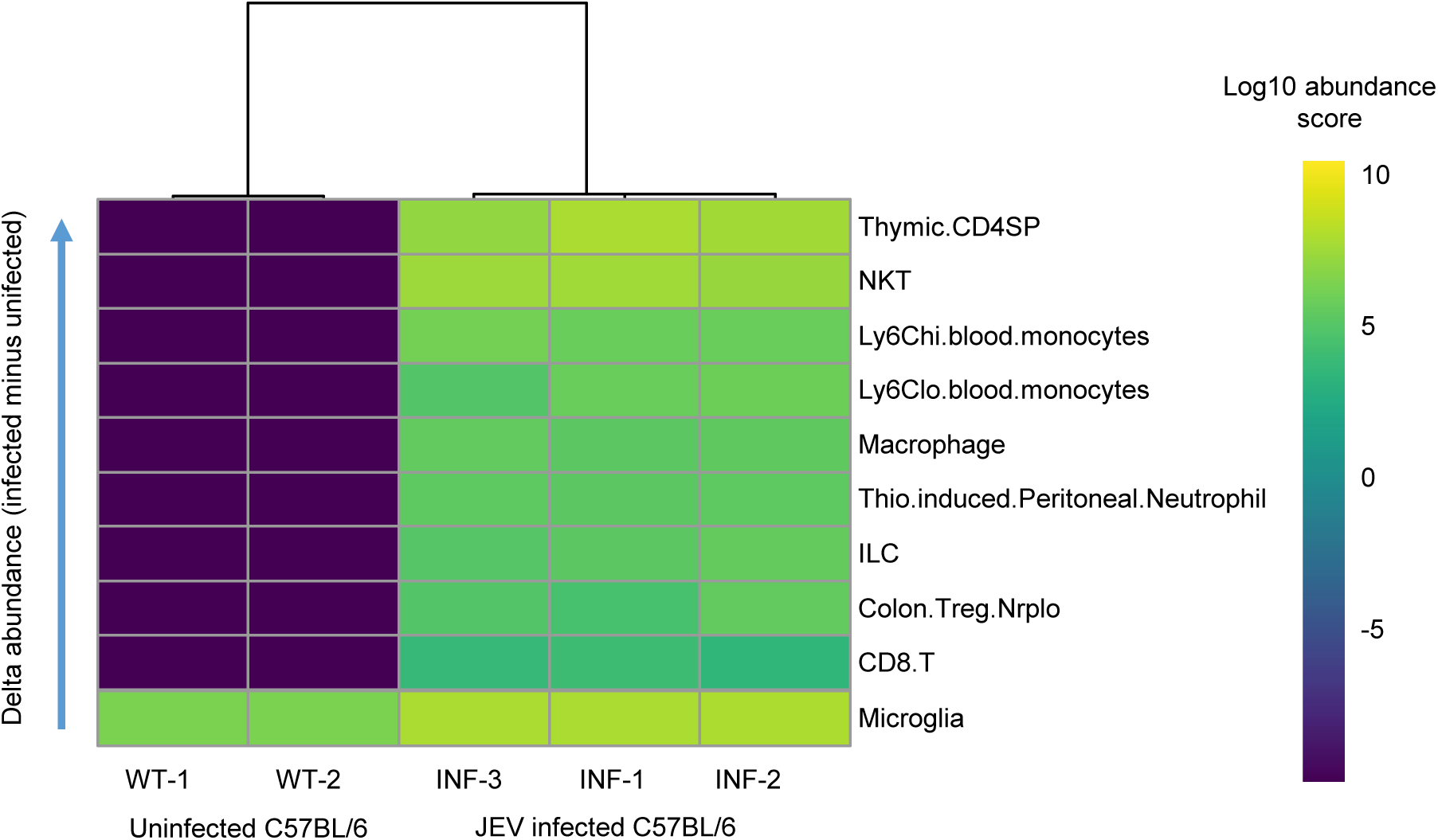
RNA-Seq read counts from brains of C57BL/6 JEV-infected and mock-infected mice obtained from the Gene Expression Omnibus (GEO accession: GSE154002) were normalised for sequencing depth and composition using DESeq2. An estimation of cell type abundances was performed on normalised read counts with the SpatialDecon package in R, using an adult mouse immune cell gene expression reference (Yoshida *et al.* 2019. Cell 176; 897-912.e20). Log_10_ abundance scores are shown for cell types that were significantly different between infected and mock-infected groups (t-test, p-value < 0.05), with cell types ordered from largest to smallest difference in mean abundance score between groups.

**Supplementary Fig. 13.**
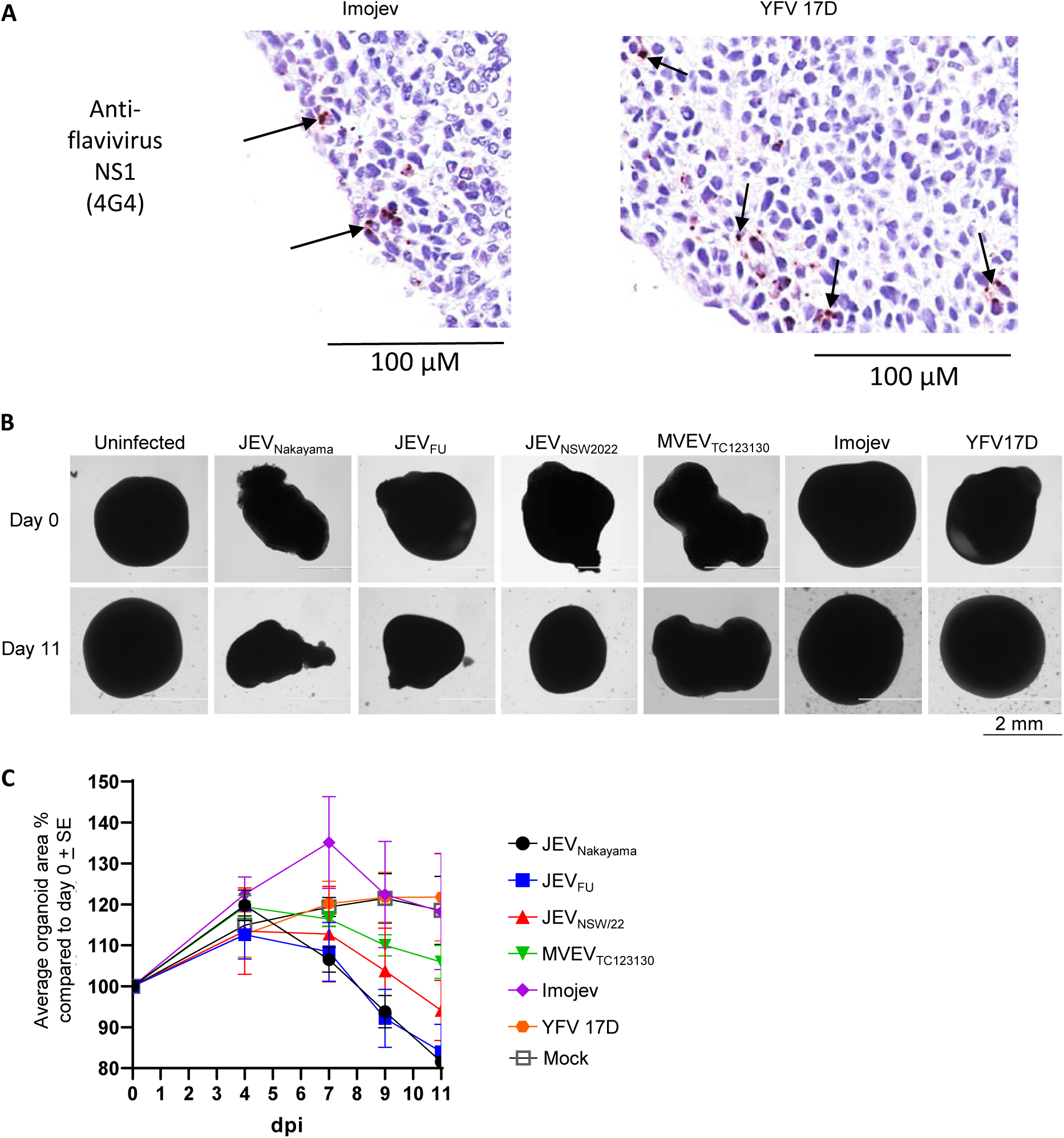
Human cortical brain organoids measurements and magnified virus staining. (A) Occasional cells staining positive for IHC for viral antigen in hBOs infected with Imojev or YFV 17D. (B) Light microscopy images of hBOs infected with JEV_Nakayama_, JEV_FU_, JEV_NSW2022_, MVEV_TC123130_, Imojev or YFV 17D at day 0 and day 11 post infection compared to uninfected hBOs. Scale bar is consistent across all images. (C) Change in hBO area on day 4, 7, 9 and 11 compared to day 0. n=8 for uninfected and MVEV_TC123130_, n=4 for all others.

**Supplementary Figure 14.**
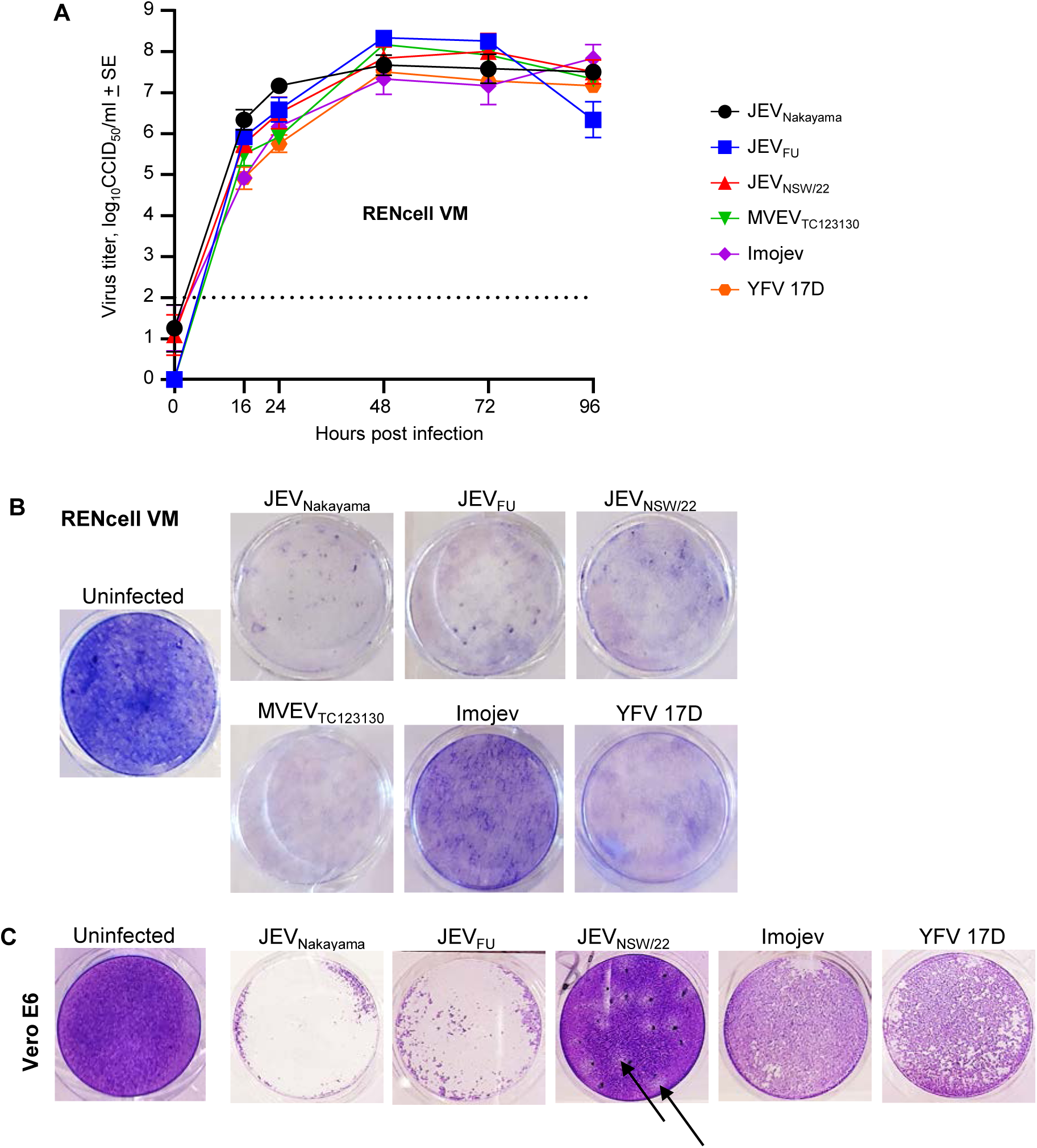
Replication and CPE in RENcell VM neural progenitor and Vero E6 cell lines. (A) Viral growth kinetics in RENcell VM determined by CCID_50_ assays of culture supernatant at the indicated times post infection. Data is the mean of 6 replicates per virus isolate across 2 independent experiments; (limit of detection is 2 log_10_CCID_50_/ml). (B) Images of crystal violet stained RENcell VM at 6 dpi (representative of n=3 per group). Less blue/violet stained cells indicates more viral CPE. (C) Vero cells were seeded at 2.5×10^5^ cells per well in 24 well plates overnight at 37°C. Cells were infected at MOI=0.05 for 1 hr at 37°C before overlay media (0.375% w/v high viscosity carboxymethyl cellulose [CMC, Sigma-Aldrich]/RPMI 1640 substituted with 2% FCS) was added to each well. Plates were incubated for 5 days before the media was removed and monolayers were fixed and stained with 0.1% w/v crystal violet (Sigma-Aldrich) in formaldehyde (1% v/v) and methanol (1% v/v). Plates were washed in tap water and dried before images were taken. Images shown are representative of n=4 per group. CPE is less pronounced in JEV_NEW/22_ infected cells, with plaques (arrows), rather than fulminant CPE, apparent at 5 dpi.

**Supplementary Fig. 15.**
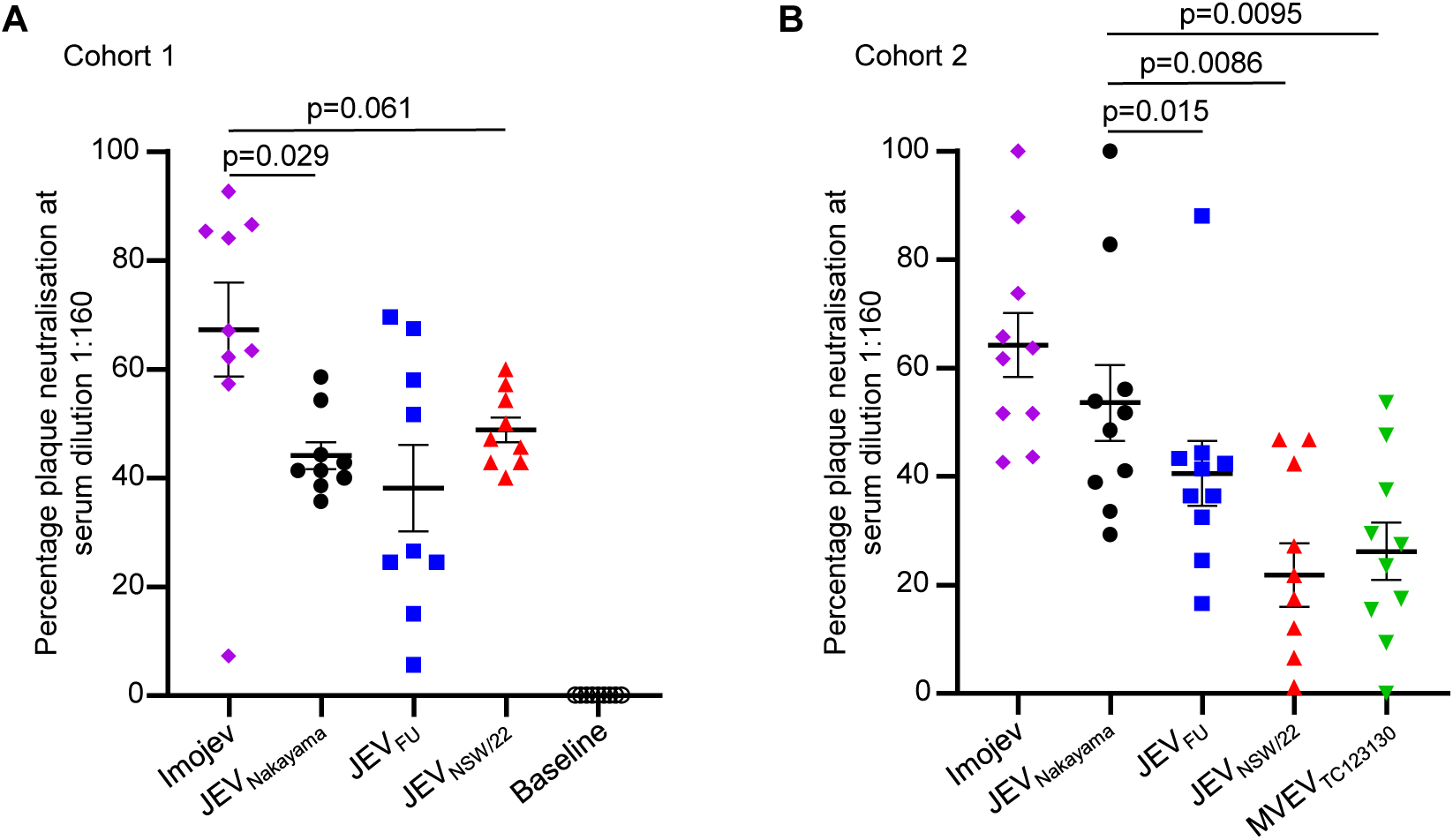
Percentage neutralisation of JEV and MVEV at serum dilution 1:160. (A) Human serum from cohort 1 (n=9) taken taken pre-vaccination (baseline) and day 28 post-Imojev vaccination was used in plaque reduction neutralisation assays against Imojev, JEV_Nakayama_, JEV_NSW/22_ and JEV_FU_. (B) Human serum from cohort 2 (n=10) taken >2 months post-Imojev vaccination was used in plaque reduction neutralisation assays against Imojev, JEV_Nakayama_, JEV_NSW/22_ and JEV_FU_ and MVEV_TC123130_. Individual data points in both A and B represent the mean percentage neutralisation at serum dilution 1:160 from duplicate wells in PRNT_50_ assay. The mean of all individuals (n=9 from cohort 1 and n=10 from cohort 2) and standard error is shown. Statistics are by paired t-test.

**Supplementary Fig. 16.**
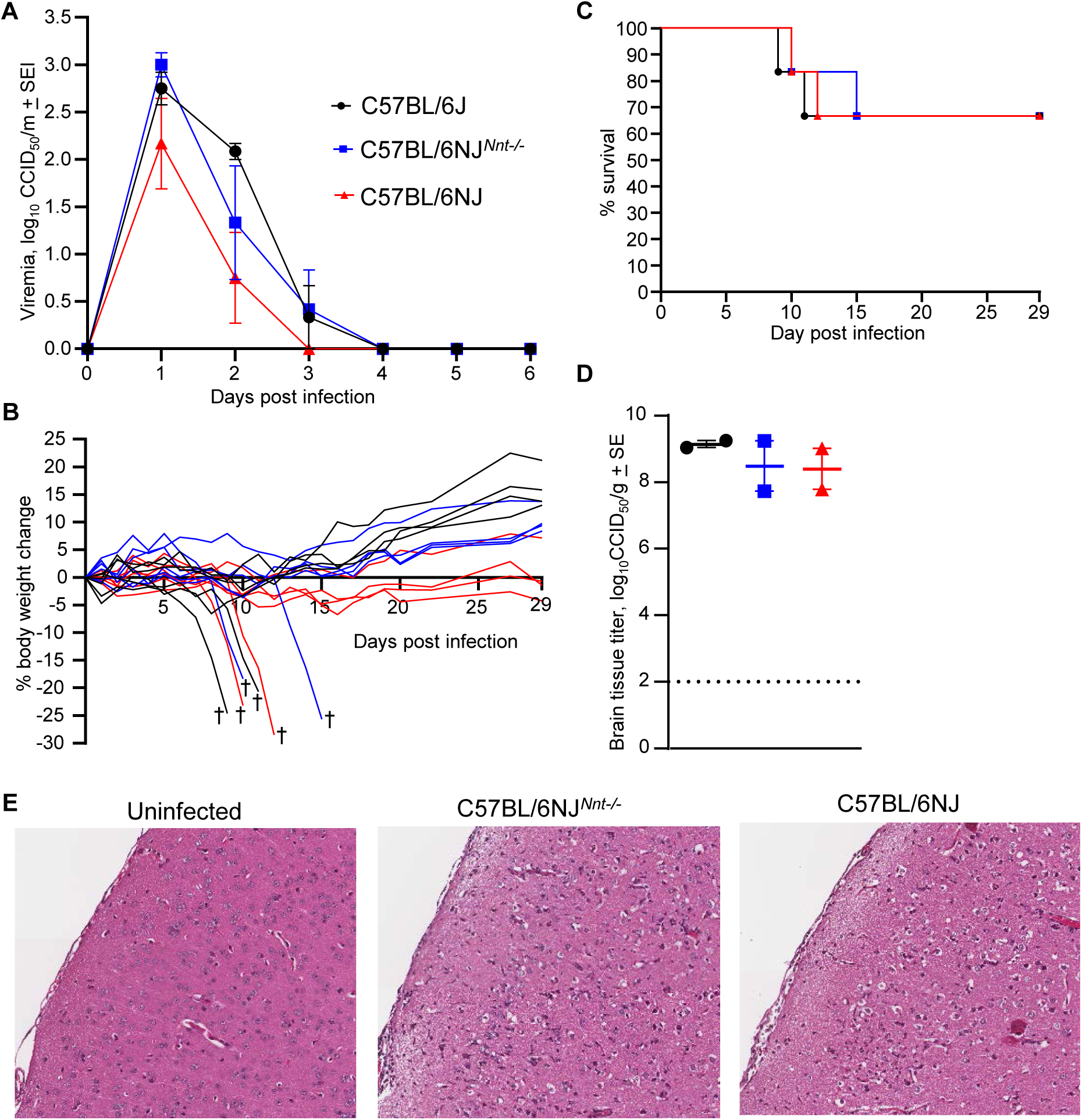
Neither mouse background nor *Nnt* significantly affected JEV neuropathogenesis. Female C57BL/6J (10-16 weeks old) (black circles), C57BL/6NJ*^Nnt-/-^* (blue squares), and C57BL/6NJ (red triangles) mice were infected s.c. with 5×10^5^ CCID_50_ JEV_Nakayama_. (A) Average viremia as determined by CCID_50_ assays; limit of detection for individual mice is 2 log_10_CCID_50_/ml. (B) Percent body weight change of individual mice compared to their weight 0 dpi. Six mice lost ≥20% body weight and required euthanasia (†). (C) Kaplan Myer plot showing percent survival (n=6 for each mouse strain). (D) Viral tissue titers in brains of the six euthanized mice (n=2 from each mouse strain). Tissue titers determined by CCID_50_ assays (limit of detection ∼2 log_10_CCID_50_/g). (E) Representative images of H&E stained sections of brains from mice that required euthanasia. No overt differences were identified between C57BL/6J, C57BL/6NJ*^Nnt-/-^,* or C57BL/6NJ mice.

